# Ceramide Connects TNF-α Signaling to Organelle Biophysical Remodeling

**DOI:** 10.1101/2025.07.26.666759

**Authors:** Ana. E. Ventura, Sandra N Pinto, Sarka Pokorna, Elad L Laviad, Abdou Rachid Thiam, Manuel Prieto, Anthony H. Futerman, Liana C Silva

## Abstract

Ceramides are essential bioactive lipids involved in cell physiology and stress responses, yet the mechanisms linking their biophysical effects on membranes to downstream cellular outcomes remain incompletely understood. Here, we demonstrate that endogenous ceramide generation in response to TNF-α-induced stress results in the rapid remodeling of the plasma membrane, triggering a cascade of cellular events. Ceramide accumulation increases membrane order and drives the formation of internalizing vesicles enriched in ceramides, which traffic through the endolysosomal system. These vesicles act as carriers that transmit membrane remodeling signals to intracellular compartments, altering their biophysical properties and function. In parallel, ceramide production reprograms lipid droplet metabolism, decreasing polarity and modifying gene expression related to lipid storage. Our findings provide direct evidence that ceramides function as biophysical integrators of the TNF-α stress response, coupling membrane order, vesicle trafficking, and lipid metabolic adaptation. This work reveals how localized lipid remodeling at the plasma membrane propagates intracellularly to support cellular adaptation and homeostasis under inflammatory stress.

## Introduction

The plasma membrane (PM) serves as a well-organized hub for signal transduction and molecular exchange, coordinating vital communication between the outside and the cell’s internal environment (1). This essential communication depends on multi-pathway network, including classical vesicular trafficking, direct physical connections through membrane contact sites (MCSs), and the specialized functions of a range of lipid-binding proteins. These mechanisms are crucial for the quick and accurate transmission of external signals, such as nutrients, growth factors, or stress responses, deep into the cell, guiding key cellular activities from gene expression and division to metabolic changes and reprogrammed metabolism (2).

Beyond vesicular transport, direct ER-PM contact sites also act as crucial structures for rapid and highly efficient non-vesicular lipid transfer (3). These junctions enable the bidirectional movement of lipids, allowing for quick and localized adjustments in membrane lipid composition and signaling in response to immediate cellular needs (4). The controlled transfer of key lipid molecules, such as cholesterol and various sphingolipids, through these contact sites and other non-vesicular pathways is crucial for maintaining membrane fluidity, organizing specialized membrane microdomains, and triggering diverse intracellular signaling pathways essential for cellular function (1, 5). This intricate, spatially and temporally controlled orchestration of signal and lipid trafficking is absolutely critical to maintaining systemic cellular homeostasis, ensuring that each organelle acquires the precise lipid composition and receives the appropriate signaling molecules required for its specialized biochemical and physiological functions (2, 4, 6).

Dysregulation within PM signaling and intracellular lipid transport pathways can have profound pathological consequences, contributing significantly to the etiology and progression of a wide spectrum of diseases, including type II diabetes, cardiovascular diseases, notably atherosclerosis, and various neurodegenerative disorders. Disruptions in sphingolipids (SL) are directly linked to the pathogenesis of lysosomal storage disorders or the progression of certain malignancies (7, 8).

Sphingolipids are essential components of eukaryotic membranes and have emerged as key regulators of cellular signaling, membrane dynamics, and metabolic homeostasis. Among them, ceramides, the hydrophobic backbone of complex SLs, are central players in stress responses, inflammation, and apoptosis (9, 10). Ceramides are generated rapidly at the PM in response to diverse stress stimuli through the action of sphingomyelinases (SMases), which hydrolyze sphingomyelin (SM) into ceramide (11–13).

Due to their extreme hydrophobicity, ceramides are unlikely to diffuse freely or act through classical soluble signaling pathways (14). Instead, it has long been proposed that their biological effects arise from the direct modulation of membrane properties. This idea is strongly supported by studies on model membranes, which demonstrate that ceramide induces significant biophysical transitions, including the formation of rigid gel domains and curvature-driven budding and tubulation (15–20). These processes are sensitive to lipid composition and ceramide species (16, 18, 20–22) and are thought to underlie the formation of “ceramide platforms”, specialized membrane domains that facilitate receptor clustering and signal transductions (13, 15). In cells, acute ceramide accumulation driven by exogenous bacterial SMase (bSMase) has been shown to induce membrane rigidification (23) and enhanced endocytosis (24, 25), further supporting the idea that ceramides can act as biophysical effectors in cellular contexts. Yet, these systems generate large, non-physiological amounts of ceramides and bypass regulatory and signaling pathways. Therefore, whether ceramide production under physiologically relevant conditions recapitulates the membrane remodeling and downstream functional changes seen in simplified systems remains unclear. Moreover, although ceramides have been linked to the activation of intracellular pathways and organelle remodeling (26), it is not known how these spatially distant effects are initiated by localized, stress-induced ceramide generation at the PM.

Here, we investigate whether endogenously generated ceramides at the PM, in response to stress, can induce functional and biophysical remodeling in human cells. Using a combination of lipidomics, live-cell imaging, and spectroscopic approaches, we show that ceramide accumulation following tumor necrosis factor-α (TNF-α) stimulation, a stress signal known to activate neutral SMase (nSMase) (27–29), triggers temporally coordinated cellular responses that link changes in membrane properties to PM remodeling and rapid endocytic internalization via ceramide-induced vesicle formation. These effects are not confined to the cell surface but extend to endosomal and lysosomal compartments, revealing that ceramide-dependent membrane remodeling operates across multiple levels of cellular organization. We further demonstrate that ceramide elevation directly affects the physical state and metabolism of lipid droplets (LDs), thereby reinforcing the link between membrane stress and lipid storage pathways (30–32).

Together, our findings demonstrate that ceramide acts as a central biophysical integrator of the cellular stress response, linking changes in membrane order and curvature at the PM to downstream pathways of endocytic trafficking and lipid storage. This coordinated, lipid-based response unfolds across multiple organelles, offering a mechanistic explanation for how ceramide exerts its widespread effects despite being confined to membranes. By coupling membrane remodeling with metabolic adaptation, ceramides may enable cells to buffer stress and reestablish homeostasis through dynamic reorganization of membrane architecture and lipid handling.

## Results

### TNF-α stimulation remodels sphingolipid metabolism and promotes membrane ordering through ceramide formation

TNF-α stimulation induced internalization of the TNF receptor in HEK cells, as visualized by confocal microscopy (Fig. 1A), without causing significant cytotoxicity (Fig. 1B). Enzymatic assays revealed a rapid and selective activation of neutral sphingomyelinase (nSMase), with no change in acid sphingomyelinase (aSMase) activity (Fig. 1C). This was accompanied by progressive hydrolysis of sphingomyelin (SM) (Fig. 1D) and ceramide formation, indicative of nSMase-dependent lipid remodeling.

**Figure 1.**
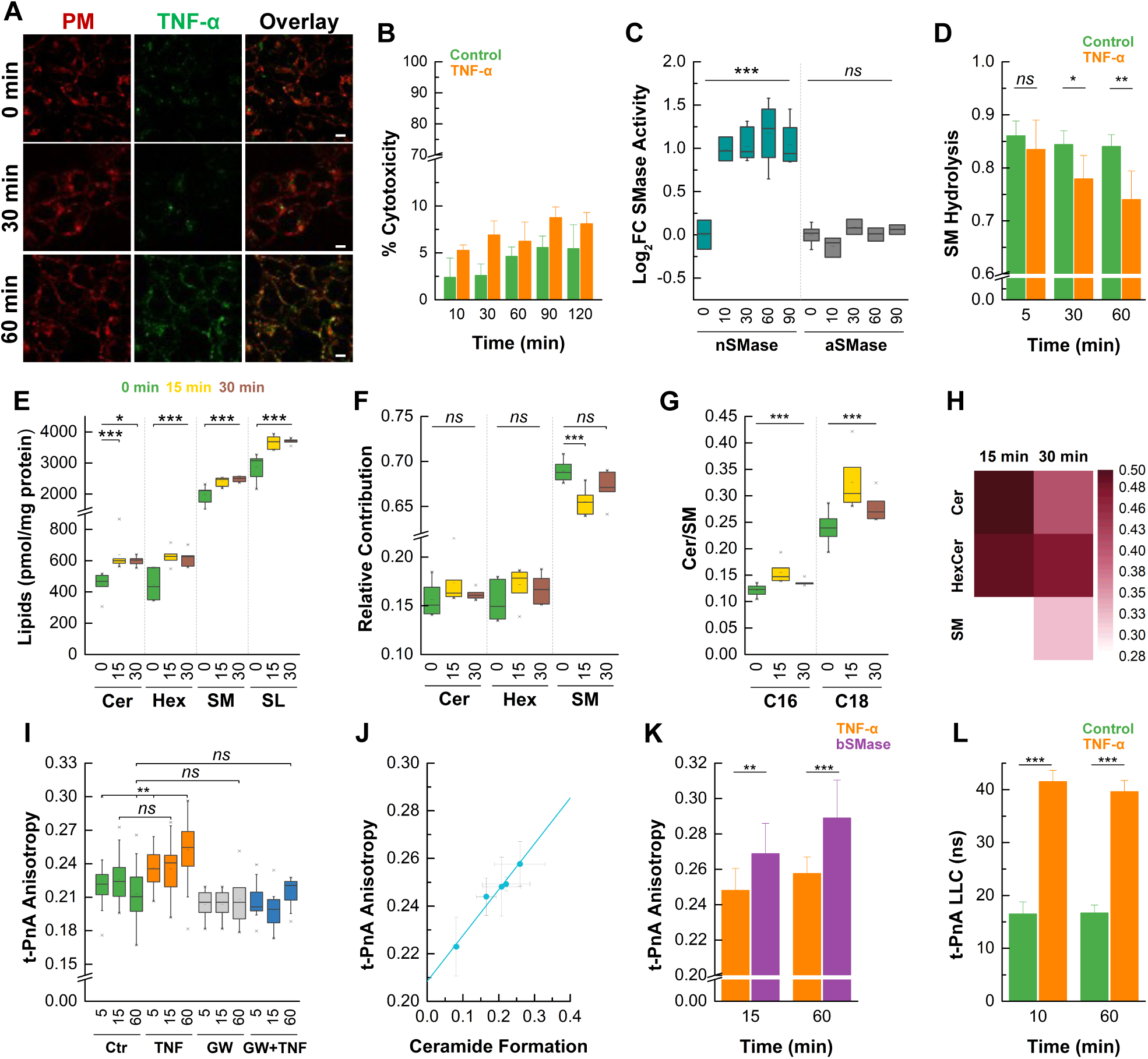
Activation of TNF receptor causes changes in sphingolipid metabolism and membrane fluidity. **A)** HEK cells were stimulated with a complex of TNF-α-biotin-Avidin-FITC and internalization of the receptor (green) was followed overtime by confocal microscopy. Cells were also labelled with WGA-Alexa 594 (red) for 10 min for PM staining. Scale bar 15 μm. **B)** Quantification of cytotoxicity (% cell death) at indicated time points after TNF-α treatment. **C)** TNF-α-induced activation of neutral (nSMase) and acid (aSMase) sphingomyelinases shown as log₂ fold change (Log₂FC) relative to control. Data show selective activation of nSMase. **D)** Quantification of SM hydrolysis indicating time-dependent reduction in SM levels upon treatment with TNF-α. **E-H)** ESI-MS/MS was used to evaluate TNF-α-induced changes in sphingolipids profiles. **E)** Quantification of the total levels of ceramide (Cer), hexosylceramide (Hex), sphingomyelin (SM) and sphingolipids (SLs) before (0 min) and 15 and 30 min after TNF-α treatment. **F)** Relative contribution of each lipid species to the sphingolipids pool, revealing a time-dependent shift in sphingolipid composition following TNF-α stimulation. **G)** Ceramide to sphingomyelin (Cer/SM) ratios for C16 and C18 species before (0 min) and 15-, and 30-min post-TNF-α treatment. The increase in Cer/SM ratio reflects a shift toward the accumulation of these ceramide species. **H)** Heatmap showing Log₂FC in levels of Cer, HexCer, and SM after TNF-α treatment at 15 and 30 min compared to untreated controls. Darker shades represent greater fold changes. **I)** t-PnA (trans-parinaric acid) fluorescence anisotropy measurements in control cells and cells treated with TNF-α, nSMase inhibitor (GW4869), or both, over time. TNF-α treatment leads to a significant increase in membrane order, which is reversed by nSMase inhibition. **J)** Linear correlation between t-PnA anisotropy and ceramide formation (data from panels D and I). **K)** A greater increase in t-PnA fluorescence anisotropy was observed in cells treated with bacterial sphingomyelinase (bSMase) compared to TNF-α, consistent with higher ceramide production and enhanced membrane rigidification. **L)** Long lifetime component (LLC) of t-PnA fluorescence intensity decay in control cells and cells treated with TNF-α. LLC values above 30 ns indicate ceramide-rich gel phase formation, consistent with increased membrane order.

Targeted lipidomics analysis revealed the rapid and coordinated remodeling of sphingolipids in response to TNF-α stimulation. A marked accumulation of ceramides (Cer) was observed at 15- and 30-min post-treatment, representing the largest fold change among the sphingolipid species (Fig 1E,H). This was accompanied by increased levels of hexosylceramides (Hex), sphingomyelins (SM), and total sphingolipids (SL), suggesting the activation of both catabolic and anabolic pathways (27). Notably, the relative contribution of sphingomyelin to the total sphingolipid pool decreased following TNF-α treatment (Fig. 1F), consistent with its enzymatic hydrolysis and conversion into ceramides. Species-level analysis revealed a selective enrichment in long-chain saturated ceramides and Hex, whereas unsaturated long-chain SM species showed the highest increase relative to control (Fig S1A-B). These changes resulted in a significant rise in the Cer/SM ratio for both C16 and C18 species (Fig. 1G, S1C). Hex/Cer ratios remained largely unchanged (Fig. S1D), indicating that glycosylation flux was not significantly altered. Principal component analysis (PCA) of the lipid profiles confirmed that TNF-α stimulation drives distinct temporal shifts in the lipidome, with clear separation between control, 15 min, and 30 min samples along PC1 (Fig S1E). Overall, these results suggest a metabolic adaptation to cell stress through the modulation of cell SL metabolism, achieved by maintaining a proper balance between the different SL species.

Because SLs are central to the regulation of organelle membrane biophysical properties and function, we tested whether altered SL levels modified membrane characteristics. Biophysical measurements using the membrane probe trans-parinaric acid (t-PnA) revealed a time-dependent increase in fluorescence anisotropy following TNF-α stimulation (Fig. 1I), indicating enhanced membrane ordering. This effect was blocked by the nSMase inhibitor GW4869 and mimicked to a higher extent by exogenous bSMase treatment (Fig. 1K) (23), highlighting the role of ceramide generation in driving membrane rigidification (Fig. S2A). A linear correlation between ceramide levels and t-PnA anisotropy (Fig. 1J, Table 1) further supports a causal relationship between sphingolipid remodeling and changes in membrane order. Moreover, TNF-α treatment increased the long lifetime component (LLC) of t-PnA fluorescence decay above 30 ns (Fig. 1L), a hallmark of ceramide-rich gel domain formation (21, 23, 33), supporting the hypothesis that ceramides form highly-ordered domains in mammalian cell membranes under these stress conditions.

**Table 1.** Linear regression parameters for relationships between experimental measurements in TNF-α–stimulated cells. Each comparison represents the relationship between two experimental measurements assessed through linear regression. The table reports the best-fit slope and intercept with their respective standard deviations (± SD), and the correlation coefficient (R²) indicating the goodness of fit. Figure labels correspond to those used throughout the manuscript.

| Figure | Comparison | Slope (± SD) | Intercept (± SD) | $R^2$ |
| --- | --- | --- | --- | --- |
| 1J | *t*-PnA Anisotropy vs Ceramide Formation | $0.19 \pm 0.01$ | $0.21 \pm 0.003$ | 0.98 |
| 2C | FM4-64 Internalization vs Ceramide Formation | $7.24 \pm 0.59$ | $-0.03 \pm 0.13$ | 0.97 |
| 2D | FM4-64 Internalization vs *t*-PnA Anisotropy | $38.59 \pm 7.05$ | $-8.11 \pm 1.76$ | 0.88 |
| 3I | NBD Internalization vs Bodipy $I_{507}/I_{613}$ | $56.97 \pm 8.39$ | $12.07 \pm 4.16$ | 0.94 |
| 4D | PCC vs Ceramide Formation (EE) | $5.43 \pm 0.76$ | $-1.04 \pm 0.17$ | 0.94 |

### Ceramide formation triggers coordinated changes in membrane biophysical properties and internalization

In model membranes, bSMase-mediated sphingomyelin hydrolysis generates ceramide-enriched domains that reduce membrane fluidity and promote vesicle formation (23). In cells, bSMase triggers rapid PM internalization, a process known as massive endocytosis (24, 25, 34). To determine whether TNF-α elicits similar ceramide-driven effects, membrane dynamics and internalization were analyzed upon stimulation (Fig. 2). TNF-α induced a rapid increase in FM4-64 internalization, reflecting enhanced endocytic activity (Fig. 2A, B, S2B). This effect was abolished by pharmacological inhibition of nSMase with GW4869 and further amplified by treatment with bSMase, supporting that ceramide production was the main driver of membrane internalization (Fig. 2B). A strong linear correlation between FM4-64 uptake and intracellular ceramide levels in TNF-α-treated cells (Fig. 2C, Table 1) further indicates that this endocytic activity is tightly regulated by ceramide generation. Moreover, FM4-64 internalization also correlated with changes in membrane order, as measured by t-PnA anisotropy (Fig. 2D, S2C, Table 1), linking trafficking dynamics to ceramide-induced biophysical remodeling of the membrane.

**Figure 2.**
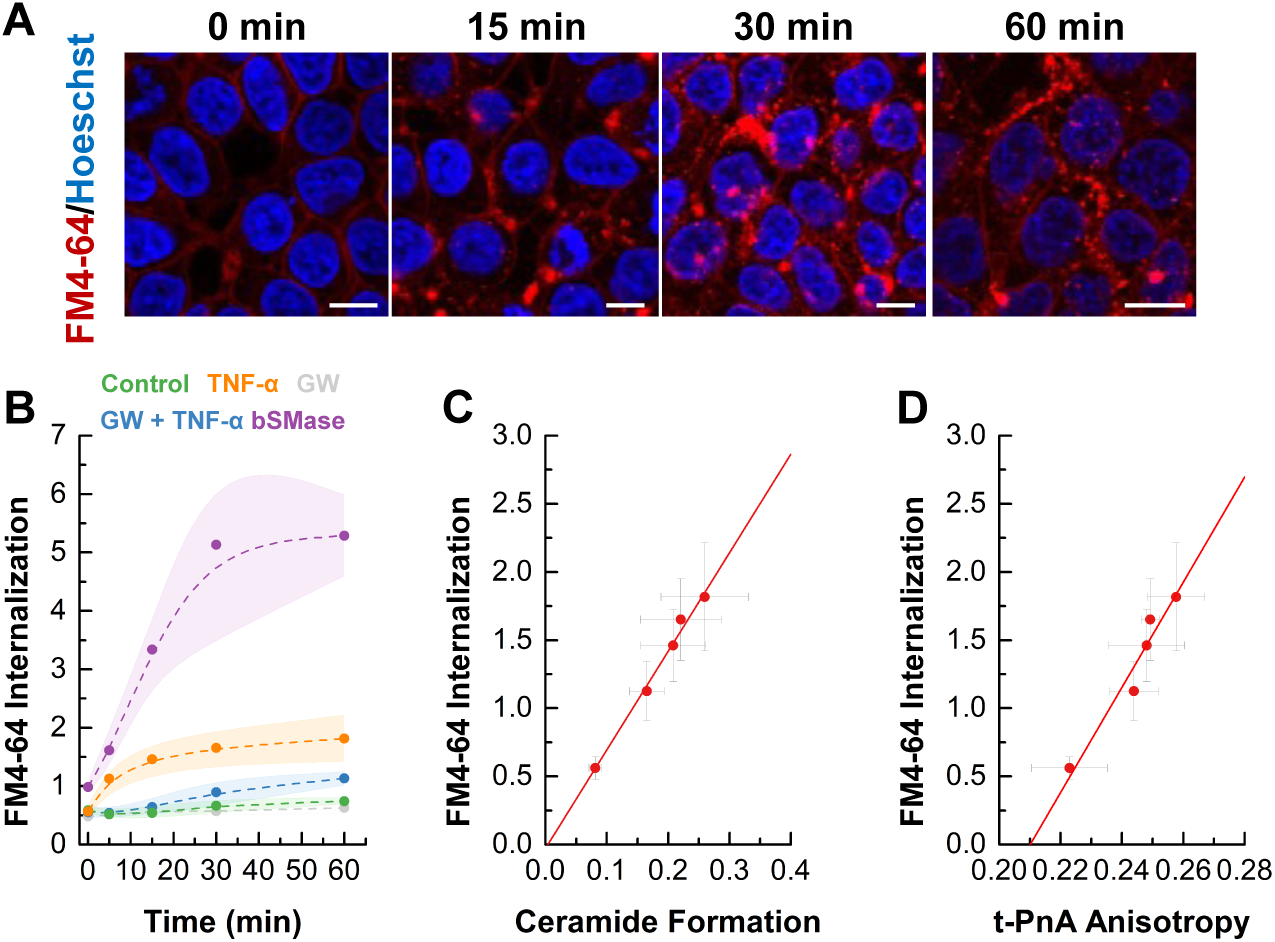
Ceramide formation causes coordinated changes in membrane fluidity and internalization. **A)** Representative confocal images of FM4-64 internalization in cells treated with TNF-α for the indicated times. Nuclei was stained with Hoechst (blue). Scale bar, 15 μm. **B)** Quantification of FM4-64 uptake over time in control cells and those treated with TNF-α, GW4869, TNF-α + GW4869, or TNF-α + bSMase. TNF-α significantly enhances internalization, which is suppressed by nSMase inhibition. Shaded areas represent SD of at least three independent experiments. Dashed lines are included solely to guide the eye and do not represent curve fitting **C, D)** FM4-64 uptake correlates linearly with **C)** ceramide levels and **D)** t-PnA anisotropy, linking endocytic activity to changes in ceramide membrane composition and biophysical state.

To investigate whether internalized membranes were enriched in ceramides, cells were labeled with the fluorescent analog C6-NBD-sphingomyelin (NBD-SM). Under basal conditions, NBD-SM localized predominantly to the PM, with minimal internalization, in contrast to C6-NBD-Cer, which rapidly accumulated at the Golgi (Fig. S3A) (35). Upon TNF-α stimulation, a progressive internalization of NBD-labeled lipid was observed, forming distinct intracellular vesicles (Fig. 3A). This process was blocked by nSMase inhibition, confirming its dependence on ceramide production (Fig. 3B). To further investigate the mechanism, phase-separated DOPC/SM/Chol GUVs labeled with NBD-SM were treated with bSMase, leading to ceramide-rich domain formation and vesiculation (Fig. 3C). These changes were accompanied by altered membrane fluidity, with ceramide-rich domains partially excluding the probe and coexisting with a phase of intermediate order, i.e., less fluid than the initial liquid disordered (*l*_d_) but more disordered than the liquid ordered (*l*_o_) (Fig. 3C).

**Figure 3.**
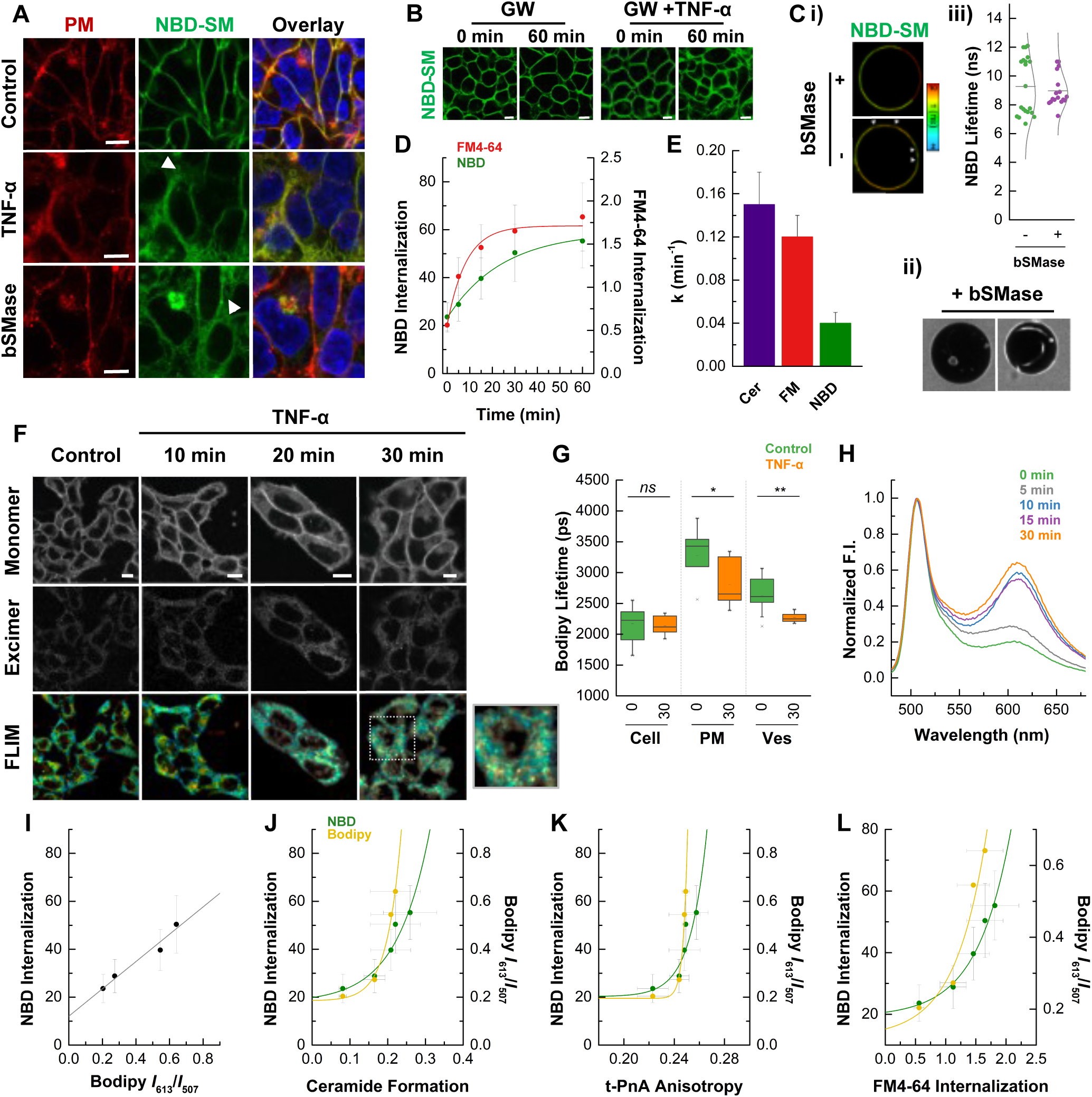
TNF-α modulates membrane dynamics and promotes internalization of ceramide-enriched vesicles. **A)** Representative confocal images of HEK cells stained with AlexaFluor 594-conjugate for the PM (red), NBD-SM (green), and Hoeschst dye (blue) in control cells and cells stimulated with TNF-α or bSMase. Arrows indicate intracellular NBD-Cer accumulation. **B)** Internalization of NBD-lipid analog was not observed in cells treated with GW4869 or GW4869+TNF-α. Scale bars, 5 μm. **C)** GUVs composed of DOPC:SM:Chol (1:1:1) exhibited clear *l*_o_/*l*_d_ phase separation, as indicated by i) NBD-SM fluorescence lifetime imaging (upper image, -bSMase). Upon treatment with bSMase, lipid reorganization occurred, leading to the formation of ceramide-rich, ordered domains from which NBD was partially excluded (+bSMase, indicated by arrows). ii) GUVs surrounded by free fluorescent dye revealed the formation of internal vesicles containing the dye (lower left image), confirming vesiculation upon bSMase treatment, as well as the emergence of tubular structures (lower right image). iii) Averaged fluorescence lifetimes of NBD-SM-labeled GUVs before and after bSMase treatment. Each dot represents an individual GUV. **D)** Time course of NBD-SM and FM4-64 internalization following TNF-α stimulation. Data points represent mean ± SD of the fluorescence intensity fraction inside the cell; solid lines indicate one-phase exponential fits, showing the kinetics of lipid/probe internalization (Table 2). **E)** Comparison of rate constants derived from one-phase exponential fits (Table 2) of ceramide accumulation, FM4-64 internalization, and NBD-SM internalization over time. **F)** Confocal images of HEK cells stained with C_5-_Bodipy-SM showing progressive internalization and membrane reorganization over time. Probe was excited at 488 nm and emission was collected at 500-595 nm (monomer) and 600 – 700 nm (excimer). Scale bars, 10 μm. FLIM images (lower row) are also shown for the same cells as the confocal images. FLIM images were acquired with a 780-nm excitation source and emission light was selected with SP 700 filter. Lifetime color code scale represents variations between 2.7 to 3.8 ns. Inset shows a representative vesicle region at higher magnification. **G)** Long lifetime component obtained for C_5-_Bodipy-SM reveals TNF-α-induced changes in lipid packing in whole cells, plasma membrane, and vesicles**. _H_)** Normalized emission spectra of C_5-_Bodipy-SM shows a time-dependent increase in the emission of Bodipy at higher wavelengths following TNF-α exposure, corresponding to increased excimer formation. **I)** NBD internalization correlates linearly with the Bodipy *I*_613_/*I*_507_ fluorescence ratio (excimer/monomer) (Table 1), indicating that increased ceramide internalization is associated with enhanced membrane packing and the formation of ceramide-enriched vesicles. **J-L)** NBD internalization and the Bodipy *I*_613_/*I*_507_ fluorescence ratio exhibit one-phase exponential relationship (Table 3) with **J)** ceramide levels, **K)** t-PnA anisotropy and **L)** FM4-64 internalization, supporting coordinated regulation of membrane order, lipid metabolism, and endocytic trafficking following TNF-α stimulation.

To dissect how ceramide modulates membrane internalization, time-course analyses of FM4-64 and NBD-SM uptake were performed in TNF-α–stimulated cells (Fig. 3D). Both probes exhibited exponential uptake kinetics over time (Table 2), but with distinct rates: FM4-64 was internalized more rapidly than NBD-SM (Fig. 3E), with a rate constant closely matching that of ceramide formation (∼1.3-fold slower). In contrast, NBD-SM uptake proceeded ∼3-fold slower than FM4-64 and ∼3.8-fold slower than ceramide, suggesting a delayed or threshold-dependent internalization mechanism. These kinetic differences indicate the presence of two ceramide-dependent endocytic pathways with distinct temporal dynamics. To further examine the relationship between probe internalization and ceramide accumulation, uptake was plotted against intracellular ceramide levels at each time point (Fig. 2C, Fig. 3J, Tables 1, 3). As noted above, FM4-64 internalization showed a linear correlation with ceramide, supporting a fast, dose-proportional uptake pathway that scales directly with TNF-α–induced ceramide production (Fig. 2C). In contrast, NBD-SM uptake followed a nonlinear (exponential) relationship with ceramide levels, reinforcing the idea of a threshold-dependent mechanism potentially involving the formation of ceramide-enriched domains and vesiculation events (Fig. 3J).

**Table 2.** Kinetic parameters from one-phase exponential fits of experimental measurements over time in TNF-α– stimulated cells. Experimental data were fit to a one-phase exponential association or decay model of the form y = A e ^(x/t)^ + y₀, where A is the amplitude, *t* the time constant, and y₀ the offset. The table presents the best-fit parameters with their associated standard deviations (± SD). The rate constant (k = 1/t) is derived from the fit. The correlation coefficient (R²) indicates the goodness of fit.

| Figure | Comparison | $y_0$ (± SD) | A (± SD) | $t$ (± SD) (min) | $k$ ($1/t$) ($\text{min}^{-1}$) | $R^2$ |
| --- | --- | --- | --- | --- | --- | --- |
| S2A | Ceramide formation | $0.23 \pm 0.01$ | $-0.15 \pm 0.01$ | $-6.62 \pm 1.54$ | $-0.15 \pm 0.03$ | 0.98 |
| S2A | *t*-PnA anisotropy | $0.25 \pm 0.001$ | $-0.03 \pm 0.01$ | $-3.36 \pm 1.98$ | $-0.30 \pm 0.17$ | 0.73 |
| 3E | FM4-64 Internalization | $1.71 \pm 0.07$ | $-1.15 \pm 0.08$ | $-8.30 \pm 1.65$ | $-0.12 \pm 0.02$ | 0.99 |
| 3E | NBD Internalization | $59.07 \pm 3.17$ | $-35.93 \pm 3.07$ | $-24.37 \pm 4.70$ | $-0.04 \pm 0.01$ | 0.99 |

Supporting this hypothesis, the extent and kinetics of NBD-SM uptake were strongly enhanced in cells treated with bSMase, which generates higher levels of ceramide (Fig. S3B-F, Table S1). These results indicate that while both internalization routes are driven by ceramide, they differ in their temporal profiles and likely reflect mechanistically distinct processes: a rapid process, tightly coupled to overall ceramide production and reflected by FM4-64 uptake, and a slower route, associated with NBD-lipid internalization, that require local ceramide accumulation, membrane reorganization and vesiculation. Importantly, membrane rigidification precedes NBD uptake, suggesting that biophysical remodeling is a prerequisite for vesicle formation (Fig. 3K, S3B).

To further confirm that these internalized membranes are enriched in ceramide and exhibit altered biophysical properties, we analyzed the structural and compositional features of the vesicles using confocal microscopy and fluorescence lifetime imaging microscopy (FLIM) with the fluorescent lipid probe C5-Bodipy-SM (Bodipy) (Fig. 3F, G). A time-dependent increase in red-shifted Bodipy fluorescence (600–700 nm) was observed in intracellular structures (Fig. 3F), consistent with excimer formation resulting from high local concentrations of the probe, indicative of ceramide clustering (36, 37). In parallel, FLIM analysis revealed a significant reduction in the fluorescence lifetime of Bodipy-lipid in both the PM and vesicles (Fig. 3G), reflecting dynamic self-quenching due to tight molecular packing and reduced probe mobility. These two effects - spectral red-shift and lifetime shortening (36, 37) - suggest that ceramide accumulation promotes local crowding of the probe within ordered lipid domains. Measurement of the emission spectra of Bodipy further confirmed this interpretation, showing a time-dependent increase in the relative intensity of the excimer peak following TNF-α stimulation (Fig. 3H).

Quantitative analysis further supports the existence of two temporally and mechanistically distinct ceramide-dependent internalization pathways. NBD-lipid internalization correlated linearly with the Bodipy excimer/monomer (*I*_613_/*I*_507_) fluorescence ratio (Fig. 3I, Table 1), indicating that both probes report on the same slower route, driven by local ceramide enrichment, organization into ceramide-enriched domains, and subsequent vesiculation. Consistent with this, and in contrast to FM4-64, both NBD and Bodipy readouts exhibited exponential relationships with ceramide levels, t-PnA anisotropy, and FM4-64 uptake (Fig. 3J–L, Table 3), supporting the idea that this delayed internalization depends on a threshold level of global ceramide accumulation and membrane remodeling. Together, these data position ceramide as a key regulator of membrane dynamics, coordinating lipid organization with vesicle formation in response to TNF-α stimulation.

**Table 3.** Kinetic parameters from one-phase exponential fits of cross-comparisons between experimental measurements. Experimental data were fit to a one-phase exponential association or decay model of the form y = A e ^(x/t)^ + y₀, where A is the amplitude, *t* the time constant, and y₀ the offset. The table presents the best-fit parameters with their associated standard deviations (± SD). The rate constant (k = 1/t) is derived from the fit. The correlation coefficient (R²) indicates the goodness of fit. Comparisons include NBD and FM4-64 internalization kinetics, t-PnA anisotropy, ceramide formation, ratio of the Bodipy fluorescence intensity at 507 and 613 nm, and Pearson correlation coefficients (PCC) obtained for early and late endosomal/lysosomal compartments.

| Figure | Comparison | $y_0 (\pm \text{SD})$ | $A (\pm \text{SD})$ | $t (\pm \text{SD})$ | $k (1/t)$ | $R^2$ |
| --- | --- | --- | --- | --- | --- | --- |
| 3J | NBD Internalization vs Ceramide Formation | $17.80 \pm 8.57$ | $2.04 \pm 3.62$ | $0.09 \pm 0.04$ | $11.42 \pm 6.35$ | 0.89 |
| 3J | Bodipy $I_{507}/I_{613}$ vs Ceramide Formation | $0.18 \pm 0.04$ | $0.001 \pm 0.002$ | $0.04 \pm 0.01$ | $26.72 \pm 7.76$ | 0.98 |
| 3K | NBD Internalization vs t-PnA Anisotropy | $20.25 \pm 8.65$ | $1.48\text{E}^{-7} \pm 1.99\text{E}^{-6}$ | $0.01 \pm 0.009$ | $75.19 \pm 51.01$ | 0.73 |
| 3K | Bodipy $I_{507}/I_{613}$ vs t-PnA Anisotropy | $0.19 \pm 0.03$ | $4.05\text{E}^{-33} \pm 7.01\text{E}^{-32}$ | $0.003 \pm 7.90\text{E}^{-4}$ | $296.16 \pm 69.37$ | 0.98 |
| 3L | NBD Internalization vs FM4-64 Internalization | $19.18 \pm 3.77$ | $1.47 \pm 1.39$ | $0.56 \pm 0.15$ | $1.79 \pm 0.49$ | 0.97 |
| 3L | Bodipy $I_{507}/I_{613}$ vs FM4-64 Internalization | $0.12 \pm 0.18$ | $0.02 \pm 0.06$ | $0.52 \pm 0.40$ | $1.92 \pm 1.48$ | 0.88 |
| 4B | PCC vs Time for Endosomes | $0.55 \pm 0.06$ | $-0.76 \pm 0.05$ | $-40.50 \pm 5.82$ | $-0.02 \pm 0.003$ | 1.00 |
| 4B | PCC vs Time for LE/Lys | $-0.42 \pm 0.24$ | $0.28 \pm 0.23$ | $61.29 \pm 35.95$ | $0.02 \pm 0.01$ | 0.97 |
| 4C | PCC vs NBD Internalization (EE) | $-0.36 \pm 0.18$ | $0.08 \pm 0.09$ | $24.40 \pm 11.12$ | $0.04 \pm 0.02$ | 0.98 |
| 4C | PCC vs NBD Internalization (LE/Lys) | $-0.13 \pm 7.92\text{E}^{-4}$ | $3.58\text{E}^{-6} \pm 5.79\text{E}^{-7}$ | $4.72 \pm 0.07$ | $0.21 \pm 0.003$ | 1.00 |
| 4D | PCC vs Ceramide Formation (LE/Lys) | $-0.18 \pm 0.07$ | $0.001 \pm 0.003$ | $0.04 \pm 0.02$ | $23.52 \pm 12.01$ | 0.89 |

### Ceramide-dependent internalization engages canonical endolysosomal trafficking and remodels lysosomal membranes

To determine whether internalized ceramide is trafficked through the endolysosomal system, colocalization analyses were performed using NBD-SM and markers for early endosomes (Rab5-RFP) and lysosomes (Lamp1-RFP) (Fig. 4A). TNF-α stimulation led to a progressive increase in colocalization of NBD signal with Rab5, followed by Lamp1 over time (Fig. 4B). Notably, the kinetic profiles of the two markers displayed opposite curvatures: Rab5 colocalization increased rapidly and approached saturation, consistent with early and transient engagement with endocytic vesicles; whereas Lamp1 association exhibited a slower onset but continued to rise over time, suggesting gradual maturation and trafficking toward late endosomal or lysosomal compartments (Table 3). These kinetics were further confirmed in bSMase-treated cells, where both Rab5 and Lamp1 colocalization with NBD increased more rapidly and to a greater extent than with TNF-α, consistent with greater ceramide production and enhanced vesicle formation (Fig. S4B, Table S1). Importantly, GW4869 treatment abolished colocalization with both markers (Fig. S4A), reinforcing the requirement for ceramide generation in vesicle formation and trafficking.

**Figure 4.**
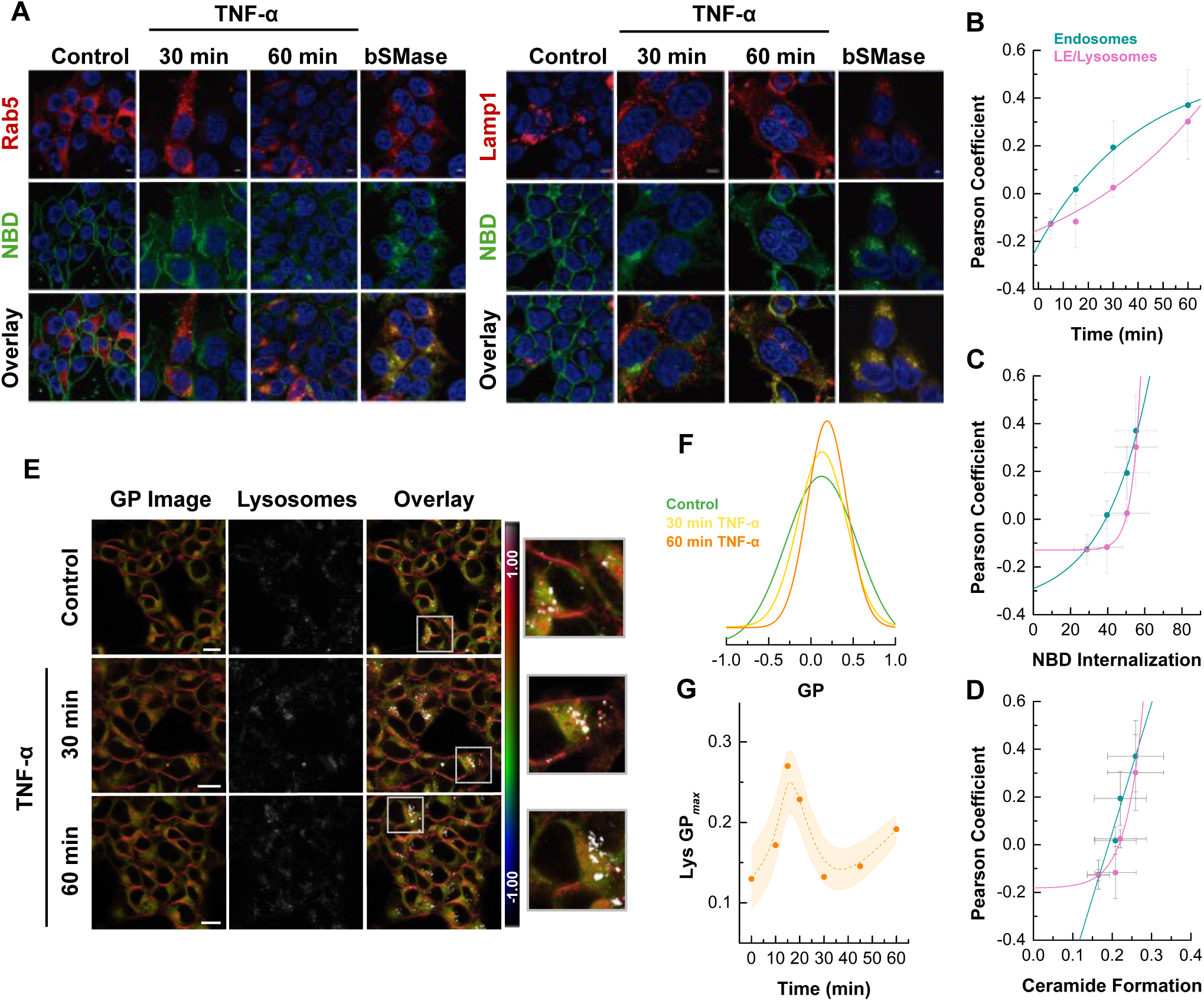
Internalized ceramide traffics to endosomal and lysosomal compartments following TNF-α stimulation. **A)** Confocal colocalization analysis of NBD-SM (green) with early endosome marker Rab5 (CellLight® Rab5-RFP, left panels) and lysosome marker Lamp1 (CellLight® Lamp1-RFP, right panels) in cells treated with TNF-α or bSMase. Merged channels show progressive accumulation of NBD signal in endolysosomal compartments in treated cells. Scale bars, 5 μm. **B)** Quantification of Pearson correlation coefficients between NBD and Rab5 (early endosomes, cyan) or Lamp1 (late endosomes/lysosomes, magenta) over time following TNF-α treatment. Data show an increasing association of internalized ceramide with both endosomal and lysosomal compartments, supporting progressive vesicular trafficking. **C, D)** Pearson correlation coefficients of NBD with early endosome and lysosome markers were plotted against **C)** NBD internalization levels measured independently (green dataset from Fig. 3D) and **D)** ceramide formation. This cross-experimental analysis links the extent of NBD internalization to its subcellular localization and ceramide formation, supporting progressive trafficking into endolysosomal compartments. Solid lines in **(B–D)** represent linear (Table 1) or one-phase exponential fits (Table 3), modeling the kinetics of colocalization and internalization over time and their relationship across conditions. **E)** Microscopy imaging of HEK cells labelled with Laurdan and LysoView 633 and treated with TNF-α. Images correspond to Laurdan GP color coded image (left), the lysosomal marker (center) and the overlay between the two (right). Scale bar, 15 μm. **F)** GP histograms show a rightward shift and narrowing of the distribution over time, indicating an increased membrane order and reduced heterogeneity in lysosome-associated membranes. **G)** Time-course analysis of lysosomal Laurdan GP_max_ values suggest transient alterations in lysosomal properties upon TNF-α stimulation. Shaded areas represent SD of at least three independent experiments. Dashed lines are included solely to guide the eye and do not represent curve fitting.

To quantitatively link internalization and compartmentalization, Pearson correlation coefficients between NBD and Rab5 or Lamp1 were plotted against NBD internalization levels (Fig 4C, Table 3) obtained from an independent experiment (data from Fig. 3E). These cross-experimental comparisons revealed a tight relationship between the degree of internalization and the extent of vesicle colocalization with endosomal and lysosomal compartments, highlighting the progressive maturation of internalized vesicles along the endocytic route. Interestingly, the correlation between Rab5 colocalization and ceramide levels followed a linear trend, suggesting that early endosomal targeting scales proportionally with ceramide production (Fig. 4D, Table 1). In contrast, Lamp1 colocalization exhibited an exponential relationship with ceramide (Fig. 4D, Table 3), indicating a mechanism that requires a critical level of ceramide accumulation for lysosomal delivery. This divergence supports a model in which early endosome engagement occurs rapidly and uniformly in response to ceramide-driven internalization, while trafficking to lysosomes requires additional maturation steps or cargo accumulation that amplify with higher ceramide loads.

To assess whether this trafficking leads to compositional changes in lysosomes, Laurdan generalized polarization (GP) imaging was combined with LysoView labeling (Fig. 4E) (38). Under basal conditions, lysosomes exhibited a broad range of GP values, reflecting significant heterogeneity in their properties. Upon TNF-α stimulation, Laurdan GP distributions progressively shifted toward higher values and narrowed over time (Fig. 4F), suggesting a transition toward more ordered and compositionally uniform lysosomal membranes. Quantification of lysosomal GP*_max_* revealed a transient peak in membrane order that declined at later time points (Fig. 4G), consistent with an early influx of ceramide-enriched membranes followed by dynamic remodeling. These findings indicate that vesicle trafficking alters the biophysical state of lysosomes, likely by transiently enriching them with ceramide and promoting membrane packing.

Together, the data demonstrate that ceramide-dependent internalization feeds into the canonical endolysosomal pathway, with vesicles progressively maturing from early endosomes to lysosomes. Rather than bypassing classical routes, ceramide-rich vesicles are incorporated into established trafficking systems, modulating the composition of recipient compartments. The magnitude and kinetics of this trafficking scale with ceramide levels, reinforcing its role as a key driver of both vesicle formation and intracellular routing.

### Ceramide generation promotes plasma membrane remodeling and increases lipid packing heterogeneity

Laurdan GP imaging was performed to further assess how ceramide formation impacts membrane organization during TNF-α stimulation. Under basal conditions, the PM exhibits higher GP values compared to intracellular regions, consistent with a more ordered membrane environment (Fig. 5A) (39). Following TNF-α stimulation, a progressive shift in Laurdan GP distributions toward higher values was observed at the PM (Fig. 5B) and across the whole cell (Fig. 5C), indicating an overall increase in lipid order. GP*_max_* values increased over time and were further enhanced by bSMase treatment, whereas nSMase inhibition with GW4869 prevented these changes (Fig. 5A–C), supporting that ceramide formation drives the observed alterations. These effects were not restricted to HEK cells. Quantification of Laurdan PM GP*_max_* in various cells revealed a consistent increase following TNF-α stimulation (Fig S5A–B), supporting that ceramide-driven membrane ordering at the PM is a generalizable feature of the TNF-α response across different cell types.

**Figure 5.**
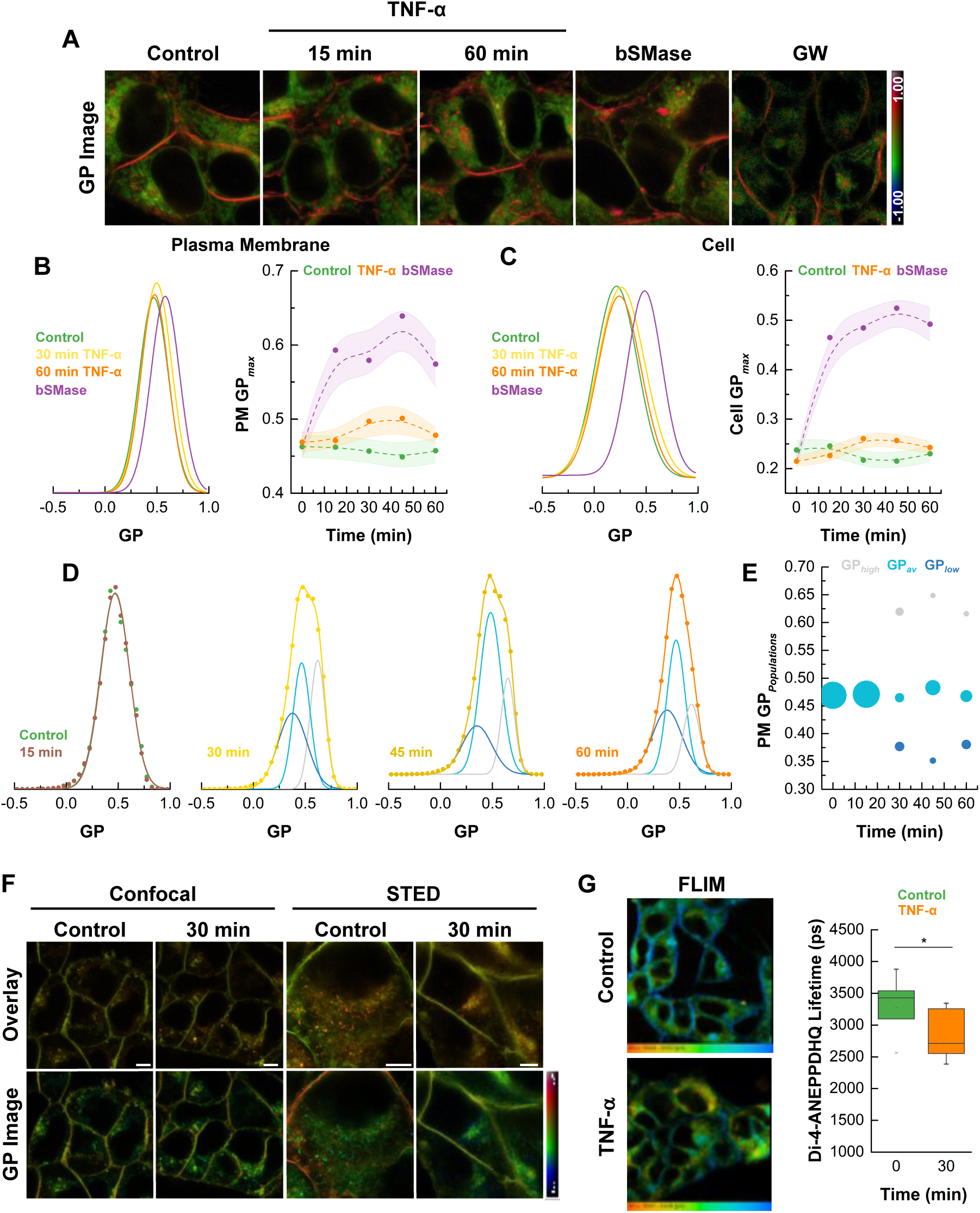
TNF-α alters membrane domain architecture and increases lipid packing heterogeneity. **A)** Representative Laurdan GP color coded images for control cells and cells treated with TNF-α, bSMase or GW4869. Images were acquired using 405 nm excitation at 37° C and the GP determined from the images collected at each Laurdan emission channel (ordered: 415 to 460 nm and disordered: 470 to 530 nm channels). **B)** Left: Distribution of PM Laurdan GP values reveals a rightward shift following TNF-α or bSMase treatment, consistent with increased membrane order. Right: Time course of PM Laurdan GP_max_ following TNF-α or exogenous bSMase treatment shows a progressive increase under both conditions. **C)** GP distribution (left) and (right) GP_max_ were also determined for the whole cell following TNF-α or exogenous bSMase, indicating global reorganization of membrane lipid packing. **D)** Laurdan GP value distributions at the PM following TNF-α treatment exhibit a progressive rightward shift and increasing heterogeneity. The emergence of multiple GP peaks suggests the formation of membrane subdomains with distinct lipid packing properties, consistent with ceramide-driven remodeling. **E)** Bubble plot representation of PM Laurdan GP populations. Each bubble corresponds to a distinct GP population identified by peak fitting of the GP distribution curves. Color encodes the GP_max_ of each population, reflecting the degree of membrane order, and bubble size reflects the fractional contribution of each population at the indicated time points following TNF-α treatment. The dynamic shift in population structure reveals an increasing prevalence of both high-GP (ceramide-enriched, ordered) and low-GP (cholesterol-depleted, disordered) membrane domains, indicating the emergence of membrane heterogeneity in response to TNF-α stimulation. **F)** Representative confocal and super resolution (STED) images of Di-4-ANEPPDHQ in untreated (control) and TNF-α-stimulated HEK cells. Overlay between the ordered (green) and disordered (red) channel and the GP color coded images are shown in the, respectively, upper and lower panels. Scale bars, 15 µm for confocal and 5 µm for STED. **G)** Left: FLIM images of Di-4-ANEPPDHQ in control and TNF-α-treated cells reveal a decrease in fluorescence lifetime. Right: Quantification of Di-4-ANEPPDHQ lifetime confirms significantly reduced PM lifetimes after TNF-α treatment, consistent with decreased cholesterol content.

In addition to changes in average GP, TNF-α stimulation led to a broadening of the GP distribution and the emergence of multiple peaks (Fig. 5D). Analysis of PM GP populations by peak fitting revealed distinct membrane environments with increasing contributions from both highly ordered (GP*_high_*) and more disordered (GP*_low_*) domains (Fig. 5E). This redistribution of membrane populations suggests that sphingomyelin hydrolysis disrupts pre-existing SM/cholesterol-rich lipid rafts. In contrast, the subsequent formation of ceramide-rich domains promotes the emergence of highly ordered, tightly packed regions that exclude cholesterol (16, 18, 20–22, 40). The resulting segregation may drive cholesterol relocation to internal membranes (18, 35, 41, 42) contributing to the appearance of low-GP, disordered membranes and potentially facilitating the rapid internalization pathway observed upon TNF-α stimulation. Consistently, the increase in GP values within internal membranes (Fig. 5C) further supports the idea of cholesterol redistribution from the PM to intracellular compartments.

To gain deeper mechanistic insight into these changes, cells were labeled with the polarity-sensitive probe Di-4-ANEPPDHQ (Di-4). Like Laurdan, Di-4 undergoes spectral shifts based on its environment, but through distinct photophysical mechanisms (43), allowing the two probes to provide complementary information on membrane remodeling. Since Di-4 is more sensitive to cholesterol content (43), it can be used to monitor alterations in cholesterol distribution. Confocal and super-resolution STED imaging of Di-4 revealed a marked decrease in GP values at the PM following TNF-α stimulation, indicating a reduction in cholesterol content and/or increased lipid disorder (Fig. 5F). These findings were corroborated by FLIM, which showed a significant decrease in Di-4 lifetime at the PM of TNF-α-treated cells (Fig. 5G), consistent with a more disordered, cholesterol-depleted PM environment. Notably, the observed Di-4 lifetime closely mirrors that reported for HEK cells following cholesterol extraction with methyl-β-cyclodextrin, further supporting the conclusion that TNF-α-generated ceramide disrupts membrane cholesterol homeostasis (44). Further support for this interpretation came from STED imaging of Di-4–labeled HeLa and SH-SY5Y cells, which revealed a decrease in PM GP values alongside a striking increase in intracellular GP values (Fig. S5C). This redistribution suggests that cholesterol was internalized or relocalized following ceramide accumulation (31), contributing to the formation of more ordered internal membranes.

Together, the data indicates that TNF-α-induced ceramide production promotes widespread and coordinated remodeling of membrane architecture. This process involves both increased membrane order at the PM and enhanced heterogeneity through the formation of distinct lipid domains, as well as a redistribution of cholesterol from the cell surface to internal compartments.

### Ceramide formation alters Lipid Droplet metabolism and properties

TNF-α stimulation or bSMase treatment led to the appearance of numerous bright, intracellular Laurdan-positive structures with high GP values, indicative of highly hydrophobic environments (Fig. 5A, 6A, and S6) (39). These structures were identified as LD based on their distinct GP profile and confirmed by co-labeling with established LD markers, such as Nile Red and LipidSpot (Fig. S6A). The GP distribution of LDs progressively shifted to higher values following TNF-α or bSMase treatment (Fig. 6B), with LD GP*_max_* values increasing over time (Fig. 6C), consistent with decreased polarity (enhanced hydrophobicity) within the LD core. This likely reflects altered LD composition, such as increased cholesteryl ester (CE) accumulation (39), driven by ceramide formation. Consistent with this, a recent study showed that TNF-α stimulation in HeLa cells promotes the formation of nuclear envelope-associated LD (NE-LD) that are highly enriched in CE, suggesting that inflammatory signaling leads to spatially regulated lipid storage remodeling (31). We further showed that LD accumulation was dependent on sphingomyelin hydrolysis, as treatment with the nSMase inhibitor GW4869 prevented the increase in LD number (Fig. 5A, S6B). The inhibition of de novo lipid synthesis with Triacsin C (45) also blocked the rise in LD GP values and reduced LD abundance (Fig. S6C, D), indicating that both ceramide production and neutral lipid biosynthesis are required for these changes. These findings suggest that ceramide generation at the PM initiates signaling and metabolic responses that regulate LD biogenesis and composition.

**Figure 6.**
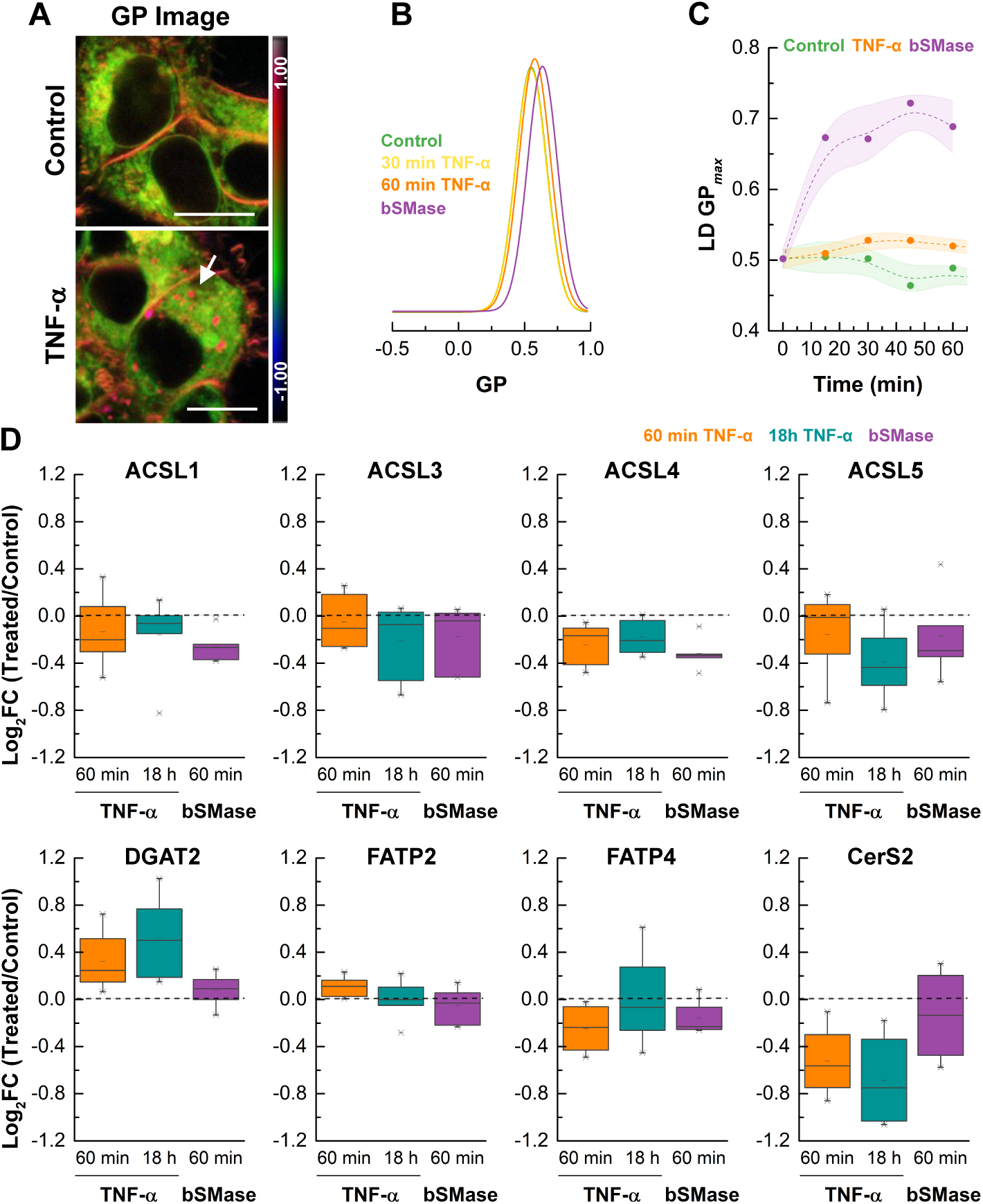
TNF-α-induced ceramide formation affects LDs metabolism and properties. **A)** Representative Laurdan GP color coded images of control cells and cells treated with TNF-α. Arrows indicate LDs with increased hydrophobicity. Scale bar, 15 μm. **B)** LD Laurdan GP distribution demonstrates a rightward shift upon TNF-α or bSMase treatment, consistent with increased LD hydrophobicity. **C)** LD GPmax increases over time both upon TNF-α or bSMase, consistent with ceramide-dependent changes. Data were collected from at least five independent experiments. More than 50 cells were analyzed for each condition. **D)** Gene expression analysis by RT-qPCR showing log₂ fold changes (treated/control) in transcripts related to lipid metabolism: ACSL1, ACSL3, ACSL4, ACSL5 (long-chain fatty acid-CoA synthetases); DGAT2 (diacylglycerol O-acyltransferase); FATP2, FATP4 (fatty acid transport proteins); and CerS2 (ceramide synthase 2). Changes were measured at 60 min and 18 h post-TNF-α or 60 min post-bSMase treatment. TNF-α significantly upregulates DGAT2 and ACSL1, potentially contributing to altered LD properties.

This ceramide-dependent remodeling of LD was not restricted to HEK cells. TNF-α stimulation also altered the properties of high-GP Laurdan-positive intracellular structures across multiple cell lines, including A549, HeLa, human macrophages, and MDA-MB231 cells (Fig. S5A, S6E). In all models, quantitative analysis revealed elevated LD GP*_max_* values, indicating a conserved increase in LD hydrophobicity following stimulation (Fig. S6E). Notably, the relative increase in LD GP*_max_* (treated/control) was more pronounced than the corresponding changes at the PM (Fig. S6F), suggesting that ceramide-driven remodeling exerts a particularly strong effect on LD core properties. These results highlight the robustness of this response across diverse cell types, underscoring ceramide’s central role in regulating LD abundance and composition.

To assess how ceramide formation influences lipid metabolic pathways, the expression of genes involved in fatty acid activation, transport, and storage was measured by qPCR (Fig. 6D). Multiple ACSL isoforms were downregulated after TNF-α or bSMase treatment, with ACSL4 showing the most potent suppression at 1 h and ACSL5 at 18 h post-TNF-α, suggesting reduced fatty acid activation in response to ceramide accumulation. DGAT2, which catalyzes triacylglycerol synthesis and promotes LD expansion, was upregulated following TNF-α treatment, with a more pronounced increase at 18 h. FATP2 and FATP4, which are involved in fatty acid uptake, showed modest or variable changes across the treatments. CerS2, which encodes a ceramide synthase for very long-chain species, was downregulated by TNF-α, suggesting a potential shift in ceramide species composition during inflammatory stress. This transcriptional suppression may contribute to altered lipid remodeling and LD phenotype.

Our results demonstrate that ceramide generation, whether through TNF-α signaling or exogenous SMase activity, modifies LD properties by promoting their formation and increasing their hydrophobicity. However, TNF-α and bSMase induce partially distinct changes in the expression of lipid metabolism genes, suggesting differential regulation of lipid handling depending on the upstream stimulus. These findings point to a broader role for ceramide in coordinating membrane remodeling and lipid storage under metabolic stress.

## Discussion

### Ceramide generation triggers membrane biophysical remodeling and vesicle internalization

Our findings reveal that TNF-α stimulation in HEK cells induces a rapid and extensive remodeling of the PM that originates from the enzymatic activation of nSMase and consequent ceramide production. Using lipidomics, we detect an increase in ceramide species, particularly C16 and C18 ceramides, as early as 15 minutes post-stimulation, coinciding with a decrease in sphingomyelin. This suggests that sphingomyelin hydrolysis is not merely a secondary response to TNF-α signaling but represents a primary biophysical trigger of downstream membrane remodeling.

Functionally, ceramide accumulation alters the physical state of the membrane, increasing lateral heterogeneity and promoting lipid reorganization. Multiple biophysical probes confirm a shift toward a more ordered and less hydrated membrane environment, consistent with ceramide-induced phase separation and the formation of tightly packed, gel-like domains. These effects mirror in vitro data showing that ceramide-rich membranes exhibit reduced fluidity, enhanced lateral segregation, and a propensity for negative curvature (46, 47). The strong linear correlation between ceramide levels and t-PnA anisotropy—a probe highly sensitive to ceramide domain formation—supports a direct link between lipid composition and membrane physical state. Notably, ceramide-induced phase separation may promote the relocalization of cholesterol from the PM to internal compartments (48), resulting in local cholesterol depletion and increased fluidity in surrounding membrane regions. This redistribution contributes to broader membrane heterogeneity and may influence the lipid composition of intracellular organelles.

This membrane remodeling initiates two kinetically distinct waves of internalization. The first, reported by FM4-64 uptake, is a rapid process that closely tracks early ceramide production and increasing membrane order. It likely reflects local sphingomyelin hydrolysis by nSMase, generating ceramide directly within the membrane. The resulting changes in curvature and order are sufficient to drive spontaneous vesicle budding, even in the absence of large-scale phase segregation. These findings are consistent with evidence from model membranes and cell-based systems showing that ceramide generation can trigger endocytosis independently of clathrin or dynamin activity (24, 25, 41).

In contrast, the second internalization phase, characterized by delayed uptake of NBD-lipid analogs and increased Bodipy-SM excimer formation, requires more extensive membrane reorganization. This likely reflects the gradual development of gel-like, ceramide-enriched domains that exclude certain lipids and serve as platforms for selective internalization. The slower kinetics of NBD-lipid uptake, along with its correlation with excimer formation, suggest that lateral segregation of ceramide precedes vesiculation and cargo sorting.

This interpretation is further supported by Laurdan GP imaging and Di-4-ANEPPDHQ FLIM reveal substantial PM reorganization following TNF-α stimulation. Laurdan detects the emergence of multiple membrane populations over time, while Di-4 FLIM shows a progressive decrease in overall lipid packing. These changes are consistent with ceramide-induced disruption of raft-like domains and cholesterol relocalization, contributing to dynamic alterations in membrane structure and function. Prior studies have shown that ceramide reduces cholesterol solubility in membranes and facilitates its internalization via vesicular and non-vesicular routes (11, 22, 25, 42, 49, 50). Altogether, the data indicate that ceramide generation triggers rapid and spatially coordinated remodeling of membrane architecture, coupling biophysical reorganization with vesicular trafficking in response to inflammatory stress.

Once internalized, ceramide-rich vesicles engage classical endolysosomal trafficking routes. We observe sequential recruitment of Rab5 and Lamp1, indicating progression from early to late endosomes. Importantly, these vesicles retain high membrane order, suggesting that ceramide-enriched membranes preserve their biophysical identity during intracellular transport. Furthermore, lysosomes in TNF-α-treated cells display a modest but consistent increase in membrane order, hinting that ceramide accumulation may extend its influence to organelle-level membrane remodeling.

Together, our data support a model in which ceramide functions as both a biochemical signal and a biophysical effector. Its production triggers an early wave of curvature-driven vesicle formation, followed by a slower phase of domain-dependent internalization and trafficking. In parallel, ceramide disrupts lipid rafts, promotes cholesterol segregation, and contributes to the long-range reorganization of membranes across multiple organelles. These processes likely coordinate lipid trafficking, storage, and adaptation during the early stages of the cellular stress response.

### Lipid storage and metabolic reprogramming follow ceramide-driven membrane remodeling

Alongside membrane remodeling and vesicle trafficking, TNF-α stimulation causes significant changes in LD metabolism and properties. We observed a notable increase in LD number and hydrophobicity. These features indicate both increased LD formation and altered lipid composition, and they strongly correlate with ceramide production. Importantly, treatment with bSMase—without TNF-α signaling—mimicked much of the LD remodeling phenotype, showing that ceramide formation alone is enough to initiate LD formation and maturation. Nonetheless, gene expression analysis revealed differential regulation of lipid metabolic enzymes in TNF-α-versus bSMase-treated cells. DGAT2 and FATP (fatty acid transport protein) were significantly upregulated following TNF-α stimulation, but not with bSMase, indicating that cytokine signaling amplifies LD biogenesis beyond ceramide generation alone (51, 52). These enzymes are involved in the activation of fatty acids and the synthesis of TAGs, and are critical for the expansion of LDs. Interestingly, recent evidence also shows that DGAT activity and TAG facilitate CE packaging within LDs (53), suggesting that its upregulation may serve to ensure efficient cholesterol storage in response to TNF-α-induced membrane remodeling. In addition, DGAT2 has been implicated in ceramide esterification and ER stress-induced LD formation (30), highlighting its multifunctional role. In contrast, ACSL4/5, enzymes involved in the activation of long-chain fatty acids, were downregulated upon treatment with TNF-α and bSMase. This reduction may reflect a feedback mechanism that limits excessive lipid flux and prevents uncontrolled neutral lipid synthesis, explaining why LD accumulation slows at later time points. Notably, CerS2, the enzyme responsible for generating very-long-chain ceramides, was downregulated in TNF-α-treated but not bSMase-treated cells. CerS2 has been shown to localize to LDs in complex with DGAT2 and ACSL5 (30), and its differential regulation suggests that distinct ceramide pools or chain-length profiles may be involved depending on the stressor.

Altogether, these molecular findings are in line with previous reports linking TNF-α stimulation to increased CE synthesis via the upregulation of ACAT1 (54) and to broader alterations in LD metabolism (54–56). Additionally, stimulation with bSMase was shown to promote cholesterol efflux from the PM and subsequent CE deposition into LDs (41, 42, 57, 58). Our results confirm and extend these observations, showing that the extent of SM hydrolysis, and hence ceramide generation, strongly influences both LD formation and internal lipid composition. The increase in LD hydrophobicity, as measured by Laurdan GP, is consistent with the enrichment of CE. Prior studies using artificial LDs have shown that Laurdan GP increases with CE-to-TAG ratios (59), supporting the idea that LDs formed under ceramide stress are CE-rich. Still, the precise nature of the lipids stored in remodeled LDs remains to be determined. They may include CE, TAG, or O-acyl ceramides (30, 41, 60), as well as possibly free cholesterol or ceramides incorporated into the monolayer. All of these species could contribute to altered LD properties. Additional support for ceramide-dependent cholesterol trafficking comes from previous reports showing that sterol analogs rapidly localize to LDs upon SMase stimulation (41), and that PM hydrolysis of SM reduces cholesterol solubility, promoting its internal redistribution (15, 25, 42, 49).

Our findings also align with recent findings describing nuclear envelope-associated LDs (NE-LDs) that accumulate CE during metabolic or autophagic stress and are closely connected to endomembrane traffic (31). These findings reinforce the idea that LDs formed during stress are not only lipid buffers, but structural hubs that coordinate membrane and metabolic responses, potentially via ER-LD or endosome–LD contact sites (32). This might represent a key axis through which ceramide-enriched vesicles intersect with lipid storage pathways.

Emerging evidence show that LD cholesterol content is a critical determinant of cellular dysfunction. A recent study demonstrated that in steatohepatitis models, cholesterol-enriched LDs in hepatocytes drive inflammation, immune recruitment, and fibrogenesis, while changes in LD composition alter gene expression and macrophage activation profiles (61). These connections underscore the physiological relevance of LD remodeling in inflammatory contexts and highlight potential links between ceramide accumulation and disease progression.

Taken together, our findings support a model in which ceramide-driven membrane remodeling not only reshapes the PM and trafficking routes, but also reprograms lipid metabolism and storage, initially by redistributing cholesterol and sphingolipids from the PM, and subsequently by activating transcriptional and biophysical changes that promote LD formation. We propose that ceramide-induced reorganization of membrane domains and LD acts to buffer excess sphingolipid and cholesterol accumulation, thereby preventing lipotoxicity and restoring membrane homeostasis during inflammatory stress. This coordinated remodeling may allow cells to rapidly adapt to environmental stress through spatially organized lipid handling and organelle cross-talk.

## Conclusions

This study provides experimental validation for the long-standing hypothesis that ceramides act as biophysical modulators of membrane organization and function. In a physiologically relevant model of TNF-α-induced stress, we demonstrate that localized sphingomyelin hydrolysis by nSMase leads to ceramide accumulation at the PM, triggering rapid changes in membrane order, curvature, and vesicle internalization. These effects propagate beyond the cell surface, influencing endosomal trafficking, lysosomal identity, and LD composition and dynamics. Importantly, these responses unfold along two coordinated axes: one driving membrane remodeling and vesicle formation, and the other reprogramming lipid storage and metabolism. By bridging PM dynamics with intracellular metabolic adaptation, our findings position ceramide not merely as a signaling lipid, but as a central integrator of stress-induced membrane and metabolic responses. This work establishes a mechanistic framework for how inflammatory signals reshape lipid architecture and cellular homeostasis across organelles.

## Methods

### Reagents

Unless otherwise specified, all reagents were obtained from commercial suppliers and used as received. C6-NBD-SM, C6-NBD-Cer, C5-Bodipy-SM FL and all the lipids were from Avanti Polar Lipids. Laurdan, t-PnA, Nile Red, Di-4-ANEPPDHQ, Wheat Germ Agglutinin (WGA)-AlexaFluor 594, WGA-AlexaFluor 633, Rab5-Alexa Fluor 594, Lamp1-Alexa Fluor 594, ER-tracker red, Mito tracker Red FM, Transferrin-Alexa 488, Phalloidin-Rhodamine and Hoechst 33342 were from Thermo Fisher. LipidSpot™ 610 and LysoView™ 633 were from Biotium. Human recombinant TNF-α and GW4869 were from Peprotech and Sigma-Aldrich, respectively. bSMase and Triacsin C were from Sigma-Aldrich.

### Cell Culture

A549, HEK293T, HeLa, HepG2, MCF-7, MDA-MB-231 and SY-SH5Y cell lines were cultured in DMEM, supplemented with 10% fetal bovine serum (Gibco), 100 U mL^-1^ of penicillin (Gibco^TM^) and 100 µg/mLof streptomycin (Gibco^TM^). HT-29 cells were grown in RPMI 1640 (Sigma-Aldrich) supplemented with 10% FBS (Gibco^TM^), 100 U/mL of penicillin (Gibco^TM^) and 100 µg/mL of streptomycin (Gibco^TM^). Cells were maintained at 37°C in a humidified 5% CO₂ incubator. Human monocyte-derived macrophages (HM) were generated by Ficoll gradient isolation followed by CD14+ magnetic sorting and differentiation for 7 days with M-CSF (20 ng/mL).

### RNA Extraction and qPCR

Total RNA was extracted using RNeasy Micro Kits (Qiagen), treated with DNase, and reverse-transcribed using qScript cDNA Synthesis Kit (Quantabio). qPCR was performed using SYBR Green PCR Master Mix (Applied Biosystems) on a 7500 Fast Real-Time PCR System (Applied Biosystems). Gene expression was normalized to *Hhprt* using the ΔCt (cycle threshold) method. Primer sequences are listed below.

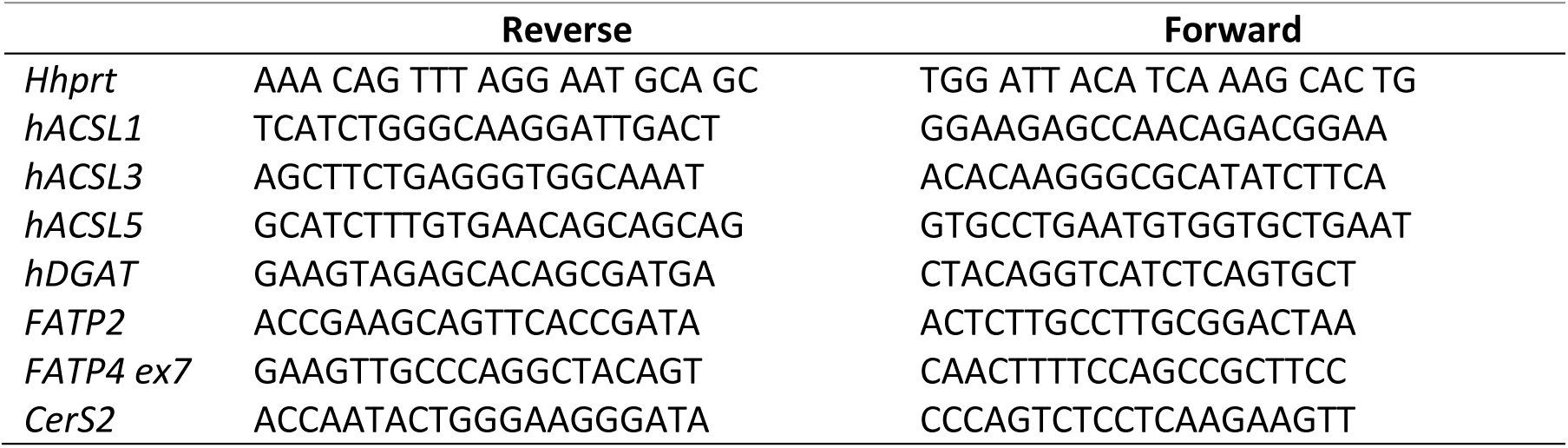

### Treatments and Pharmacological Inhibition

Cells were treated with 100 ng/mL TNF-α, 0.5 U/mL bSMase, or pretreated with 50 µM GW4869 for 30 min prior to stimulation. For inhibition of neutral lipid synthesis, cells were treated with 1 µM Triacsin C for 2 h.

### In Vitro SM-to-Cer Conversion Assay

HEK cells were labeled with 2 μM C6-NBD-SM and treated as indicated. Lipids were extracted by chloroform/methanol and analyzed by thin-layer chromatography (TLC) using C6-NBD-SM and C6-NBD-Cer standards. Fluorescent bands were imaged using a Typhoon 9410 scanner and quantified with ImageQuant TL software.

### nSMase and aSMase Activity Assay

Cells were lysed in appropriate buffer (pH 7.4 for nSMase, pH 4.5 for aSMase), incubated with 1 nmol C6-NBD-SM for 20 min at 37°C, and reactions terminated by lipid extraction. SM hydrolysis to ceramide was analyzed by TLC as above.

### Cytotoxicity Assay

LDH release was quantified using Cytotoxicity Detection KitPLUS (Roche) following treatment with TNF-α and/or GW4869. Absorbance was measured according to the manufacturer’s protocol.

### Mass Spectrometry (ESI-MS/MS)

HEK 293 cells were harvested by trypsinization, collected by centrifugation, washed twice with ice-cold phosphate-buffered saline, and lyophilized. SL analyses by ESI-MS/MS were conducted using a PE-Sciex API 3000 triple quadrupole mass spectrometer and an ABI 4000 quadrupole-linear ion trap mass spectrometer as described previously (23).

### Steady-state and time-resolved fluorescence spectroscopy

HEK cells (at ∼1×10^6^ cells/mL) were labeled with 2 μM of t-PnA or 0.5 μM C_5_-Bodipy-SM FL. Fluorescence spectra and steady-state anisotropy were measured in SLM Aminco 8100 series 2 spectrofluorimeter with double excitation and emission monochromators, MC400 (Rochester, NY) (20). All measurements were performed in 0.5 cm × 0.5 cm quartz cuvettes at 37°C using 320/405 nm for t-PnA and 500/580 and 600/650 nm for th Bodipy as excitation (λ_exc_)/emission (λ_em_) wavelengths. Time-resolved fluorescence measurements were carried out with a time-correlated single-photon timing system (62). Excitation and emission wavelength were λ_exc_ = 305 nm (secondary laser of Rhodamine 6G) and λ_em_ = 405 nm. The experimental decays were analyzed using TRFA software (Scientific Software Technologies Center, Minsk, Belarus).

### Giant Unilamellar Vesicles (GUV)

GUV were made from 18:0 sphingomyelin (SM), cholesterol (Chol) and DOPC 1,2-dioleoyl-sn-glycero-3-phosphocholine (DOPC) in ratios 10:25:65 (SM:Chol:DOPC; no phase separation) and 1:1:1 (SM:Chol:DOPC; phase separation) by electroformation described in detail previously (63). GUVs with bSMase (0.2U/mL) were imaged after 30 min of treatment.

### Cell staining

Cells were seeded in 35 mm µ-Dishes, 4-well or 8-well ibidiTreat™ µ-slide and kept in a humidified incubator at 37 °C with 5% CO_2_ for 48 h to 72 h before imaging. Cells were labeled with different dyes (2 μM C6-NBD-SM, 0.5 μM C5-SM-Bodipy FL, 0.5 μg/mL Alexa Fluor 594-conjugate, 100 μg/mL Transferrin-Alexa 488, 5µg/mL FM4-64 and 1 μg/mL Hoechst 33342 dye) and incubated at 37 °C for 10-30 min, according to supplier protocol. For co-localization studies with endolysosomal markers cells were labeled with the endosomal marker, Rab-5-RFP, or the lysosomal marker, Lamp1-RFP (CellLight® from Invitrogen), according to the manufacturer’s instructions. Cells were incubated overnight prior to imaging. On the day of the experiments, cells were labelled with 1 μM C6-NBD-SM for 10 min.

To evaluate changes on membrane biophysical properties cells were labelled with 5 μM Di-4-ANEPPDHQ (30 min), 0.5 μM C5-SM-Bodipy FL (15 min) or 5 µM of Laurdan (60 min) alone or concomitantly with 0.5 µM of NR, 1:1000 dilution of LipidSpot^TM^ 610, 1:1000 dilution of LysoView^TM^ 633, 1 μM ER-tracker red, 250 nM Mito-tracker red FM and rhodamine-phalloidin according to manufacter’s instructions. Cell medium was replaced by fresh medium without phenol red prior to imaging.

### Confocal and Multiphoton Imaging

Confocal fluorescence microscopy was performed using either a Leica TCS SP5 (Leica Microsystems CMS GmbH, Mannheim, Germany) confocal inverted microscope (DMI6000) with a 63× water (1.2 numerical aperture) apochromatic objective or a Leica TCS SP8 (Leica Microsystems CMS GmbH, Mannheim, Germany) confocal inverted microscope (DMi8 Bino) with temperature (37°C) and CO_2_ (5%) control, using a 63x water (1.2 numerical aperture) apochromatic objective. Laurdan was excited at 780 nm using a pulsed Titanium-Sapphire laser or at 405 nm using a diode laser on the Leica SP5 and Leica SP8, respectively. The emission was collected at 400-460 nm and 470-530 nm, or at 415-460 nm and 470-530 nm, when Leica SP5 and Leica SP8 were used, respectively. LipidSpot^TM^ 610 was excited using a white light laser (WLL) at 610 nm and the emission was collected at 650 – 750 nm. NR was excited at 514 nm using an Argon^+^ laser and the emission was collected at 550 – 615 nm. Cells were maintained at 37 °C, 5% CO_2_ during image acquisition in the Leica SP8 and at RT during image acquisition in the Leica SP5.

### Fluorescence Lifetime Imaging (FLIM)

FLIM experiments were carried out on a Leica SP5 inverted microscope. The microscope is equipped with a FLIM add-on comprising from Becker &Hickl. C_5_-Bodipy-SM FL fluorescence was excited at 800 nm by a Titanium:Sapphire laser and the emission decay was collected using a 560 nm short-pass dichroic bandpass filter in the descanned detection path. FLIM conditions were optimized with varying acquisition times to achieve a reasonable photon count. GUVs FLIM measurements were performed on an Olympus FluoView 1000 MPE system upgraded with a dual detector channel PicoQuant laser scanning microscope (LSM) Upgrade Kit and a home-built excitation system consisting of LDH-D-C-470 diode laser head. GUVs were imaged at room temperature. Lifetimes were determined using reconvolution procedure in SymPhoTime software (PicoQuant, Berlin, Germany).

### Super Resolution Imaging

Stimulated Emission Depletion (STED) experiments were perfomred on Leica SP8-STED inverted microscope equipped with white laser light with STED depletion lasers (594, 660 and 775). Excitation was performed at 488 nm and emission was collected at 500-580 nm and 620-750 nm.

### Data analysis and representation

Images were processed using Fiji software (Bethesda, USA). Laurdan GP was analyzed using a custom written macro for ImageJ as described in (39), and available at https://github.com/WIS-MICC-CellObservatory/LaurdanInLiveCellImaging/. The GP was calculated as described in (39) using the Laurdan images acquired in the wavelength range of 400/415-460 nm (channel 1, more ordered/hydrophobic environment) and 470-530 nm (channel 2, less ordered/more hydrophilic environment), for the 2-photon and the 405 excitations respectively. The G factor corresponds to the GP of the reference sample calculated as described in (39).

Unless otherwise indicated, data were plotted using OriginLab and presented as box-and-whisker plots showing the median, interquartile range (IQR), and full data range, or as mean ± SD of at least three independent experiments. Laurdan GP data were analyzed by extracting pixel-based GP values to generate histograms of GP distributions. These distributions were fitted with one or more Gaussian functions to resolve distinct membrane populations and characterize changes in lipid packing heterogeneity. GP*_max_* values were obtained from the mode of the fitted peaks or directly from the histogram maximum when a single peak was present. When multiple Gaussian components were required, the relative fraction of each population was calculated from the area under each peak, normalized to the total fitted area. For correlation and kinetic datasets, linear and nonlinear regression models were applied as appropriate. The type of fit used, corresponding parameters, and goodness-of-fit metrics (e.g., R²) are reported in the corresponding figure legends and summary tables (Tables 1 and 2, and Table S1).

Statistical analyses were carried out using OriginLab. Comparisons between two groups were performed using unpaired two-tailed Student’s t-tests or Mann-Whitney U tests. For comparisons involving multiple groups, one-way ANOVA followed by Tukey’s post hoc test or Kruskal-Wallis ANOVA were performed. p-values are indicated as * p < 0.05; ** p < 0.01; *** p < 0.001; *ns*, non-significant.

## Data availability statement

The manuscript has data included as electronic supplementary material and additional data will be made available on reasonable request.

## Author Contributions

L.C.S. conceptualized and supervised the study, secured funding along with A.H.F. and M.P., contributed to data visualization and administered the project. A.E.V., A.H.F. and M.P. also contributed to the conceptualization of the study. A.E.V. performed the experiments, analyzed the data, curated and visualized the results, and drafted the initial manuscript. S.N.P. and S.P. performed experiments and data analysis. E.L.L. conducted the activity assays. A.R.T., M.P., and A.H.F. contributed to data interpretation and the discussion of results. The final version of the manuscript was prepared by L.C.S. with contributions from A.R.T., A.H.F., and M.P. All authors reviewed and approved the final manuscript.

## Acknowledgements

We gratefully acknowledge Alfred H. Merrill (School of Biological Sciences & the Petit Institute for Bioengineering & Bioscience, Georgia Institute of Technology, USA) for performing the lipidomic analyses and for his valuable expertise and support throughout the study. This work was supported by national funds from FCT - Fundação para a Ciência e a Tecnologia, I.P. (Portugal) [grants PTDC/BBB-BQB/3710/2014, PTDC/BIA-BFS/29448/2017; SFRH/BD/104205/2014 to A.E.V.; UIDP/04138/2020 and SAICTPAC/0019/2015 of the Research Institute for Medicines; UIDB/04565/2020 of the Research Unit Institute for Bioengineering and Biosciences - iBB and the project LA/P/0140/2020 of the Associate Laboratory Institute for Health and Bioeconomy - i4HB]; and from the European Union’s Horizon 2020 [grant LysoMod (734825)]. S.P. acknowledges the Czech Academy of Sciences for the Czech/Israel scientific program. A. H. Futerman is The Joseph Meyerhoff Professor of Biochemistry at the Weizmann Institute of Science.

