## Supplemental data for "Ceramide Connects TNF-α Signaling to Organelle Biophysical Remodeling"

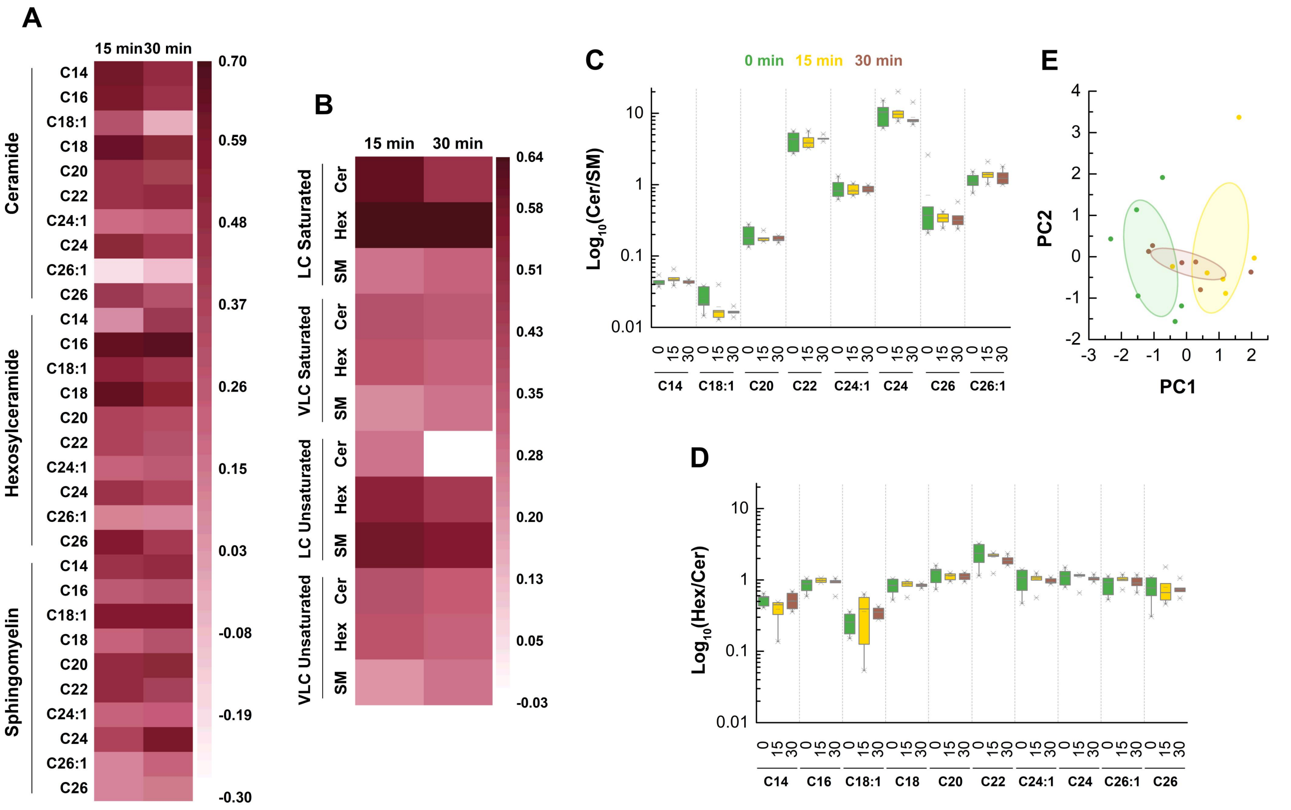


**Figure S.1. Activation of TNF receptor causes changes in sphingolipid metabolism. A)** Heatmap showing changes in individual ceramide (Cer), hexosylceramide (Hex), and sphingomyelin (SM) species following TNF-α treatment at 15 and 30 min. Values represent $Log_2FC$ compared to untreated controls. Darker shades represent greater fold changes. **B)** Heatmap of summed lipid class changes for saturated and unsaturated long-chain (LC) and very long-chain (VLC) Cer, Hex, and SM at 15 and 30 minutes. Values represent $Log_2FC$ compared to untreated controls. **C)** No significant changes were observed in the ceramide to sphingomyelin (Cer/SM) ratios for different species before (0 min) and 15-, and 30-min post-TNF-α treatment. Values represented $Log_{10}$ ratios of total levels of individual Cer/SM species. **D)** Ratio between individual Hex/Cer showing that no significant changes were observed on the glycosylation flux post-TNF-α treatment. **E)** Principal component analysis (PCA) of sphingolipid species showing temporal separation of samples based on TNF stimulation duration (0, 15, and 30 minutes). PCA reveals distinct clustering of treatment groups, with the greatest separation observed along PC1. Control and treated conditions show distinct trajectories, suggesting treatment-induced variation in lipid/metabolic profiles. Ellipses represent 95% confidence intervals for each group, based on the distribution of PC1 and PC2 scores.

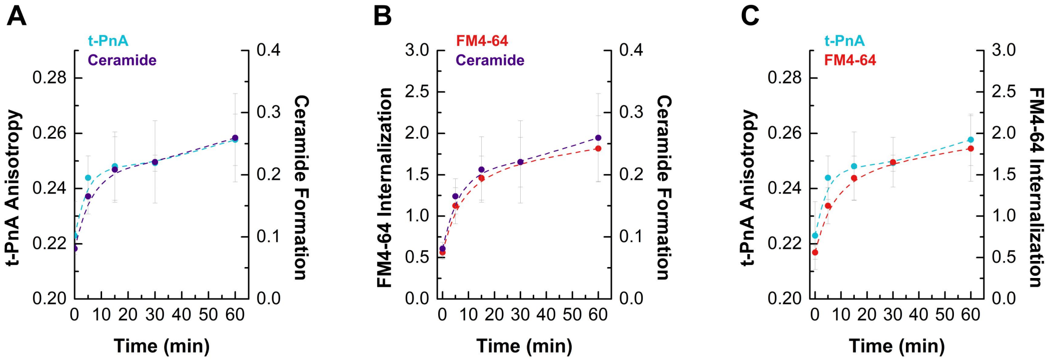


**Figure S.2. Ceramide formation causes coordinated changes in membrane fluidity and internalization. A)** Time-course of t-PnA fluorescence anisotropy showing that alterations in membrane fluidity occur in parallel with ceramide formation in HEK cells stimulated with TNF-α. **B)** FM4-64 internalization increases in correlation with rising ceramide levels in TNF-α–stimulated cells. **C)** Comparative analysis of membrane fluidity (t-PnA anisotropy) and FM4-64 internalization in HEK cells treated with TNF-α. Dashed lines are included solely to guide the eye and do not represent curve fitting. Values represent the mean ± SD of at least three independent experiments.

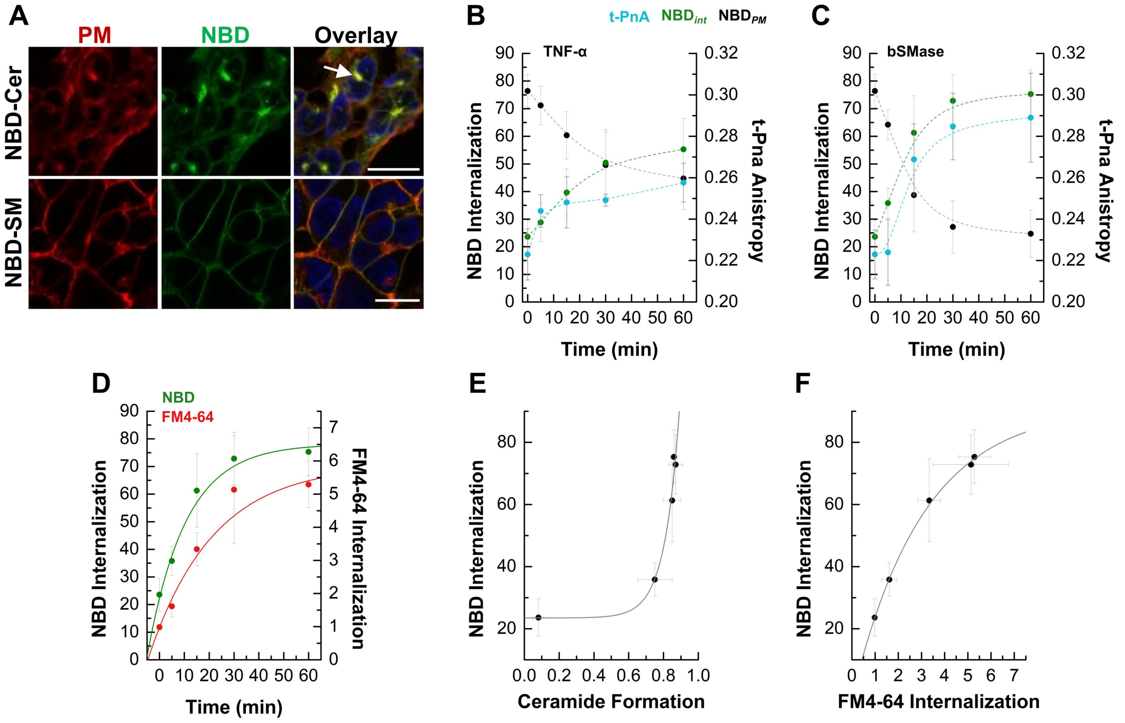


**Figure S.3. Ceramide generation correlates with membrane remodeling and internalization. A)** Untreated HEK cells labelled with (green) C6-NBD-Cer or C6-NBD-SM and stained for the PM with (red) WGA Alexa 594 showing different localization of the NBD probes: C6-NBD-Cer localizes to the Golgi while C6-NBD-SM localizes in the PM. The overlay image is also shown. Nuclei were stained with Hoescht (blue). Scale bar 15 µm. **B, C)** Total fluorescence intensity of NBD staining in the PM (black) and inside the cell (green) was determined in **B)** TNF-α and **C)** bSMase treated cells. The variation in t-PnA anisotropy is also shown (cyan). Dashed lines are included solely to guide the eye and do not represent curve fitting. **D)** Time-course of NBD and FM4-64 internalization in bSMase-treated cells. **E)** Quantification of NBD internalization as a function of ceramide formation in bSMase-treated cells. Data fitted with an exponential function reveals a nonlinear relationship (Table S1), supporting threshold-dependent uptake. **F)** Correlation between NBD and FM4-64 internalization levels in bSMase-treated cells. Data in D-F were fitted with one-exponential function (Table S1). Solid line represents the fitting.

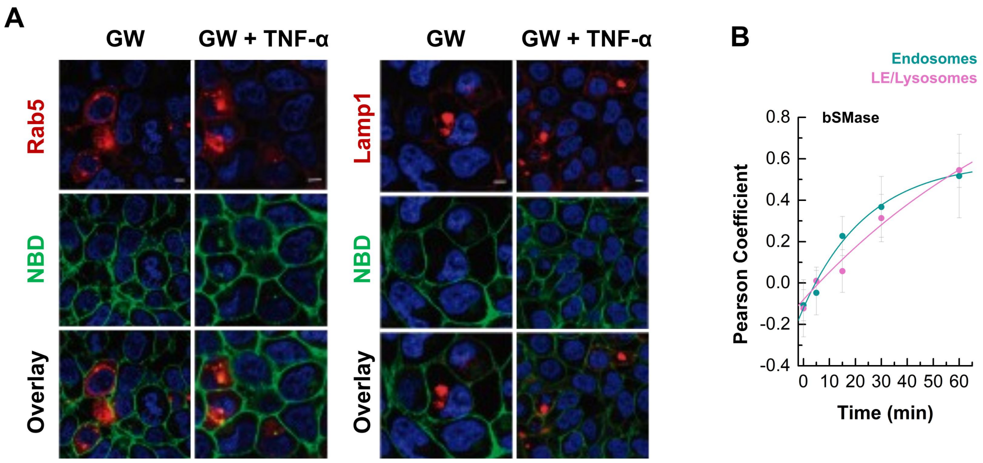


**Figure S.4. Ceramide is required for membrane internalization and trafficking to endolysosomal compartments. A)** Co-localization between (left) Rab-5-RFP and NBD (C6-NBD-SM) and between (right) Lamp1-RFP and NBD was evaluated over time upon treating cells with nSMase inhibitor with or without TNF-α. Nuclei were stained with Hoescht (blue). Scale bars, 10 µm. **B)** Quantification of Pearson correlation coefficients between NBD and Rab5 (early endosomes, cyan) or Lamp1 (late endosomes/lysosomes, magenta) over time following bSMase treatment. Fitting parameters are shown in Table S1.

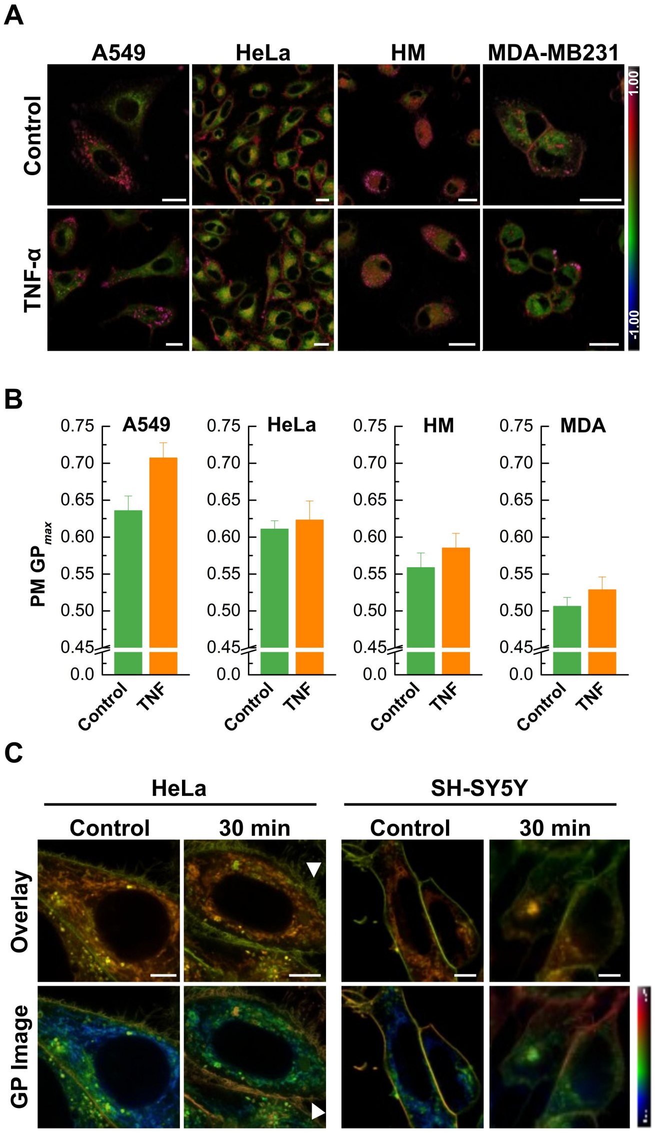


**Figure S.5. Ceramide generation increases membrane order in both plasma membrane and intracellular compartments across cell types. A)** Laurdan GP color coded images of the effect of TNF-α treatment in A549, HeLa, Human macrophages (HM) and MDA-MB231 cells. The cells were labeled with 5 μM Laurdan for 30 min and then treated with TNF-α for 30 or 60 min. **B)** Laurdan $GP_{max}$ values were determined for the PM. **C)** Control and TNF-α treated HeLa and SH-SY5Y cells were labelled for 30 min with 5 μM Di-4-ANEPPDHQ and super resolution STED images were acquired. Excitation was performed at 488 nm and the emission was collected at 500 - 580 nm (ordered channel, channel 1) and 620 - 750 nm (disordered channel, channel 2). Overlay between the ordered (green) and disordered (red) channel and the GP color coded images are shown in the, respectively, upper and lower panels. Scale bar 15 μm for confocal and 5 μm for STED.

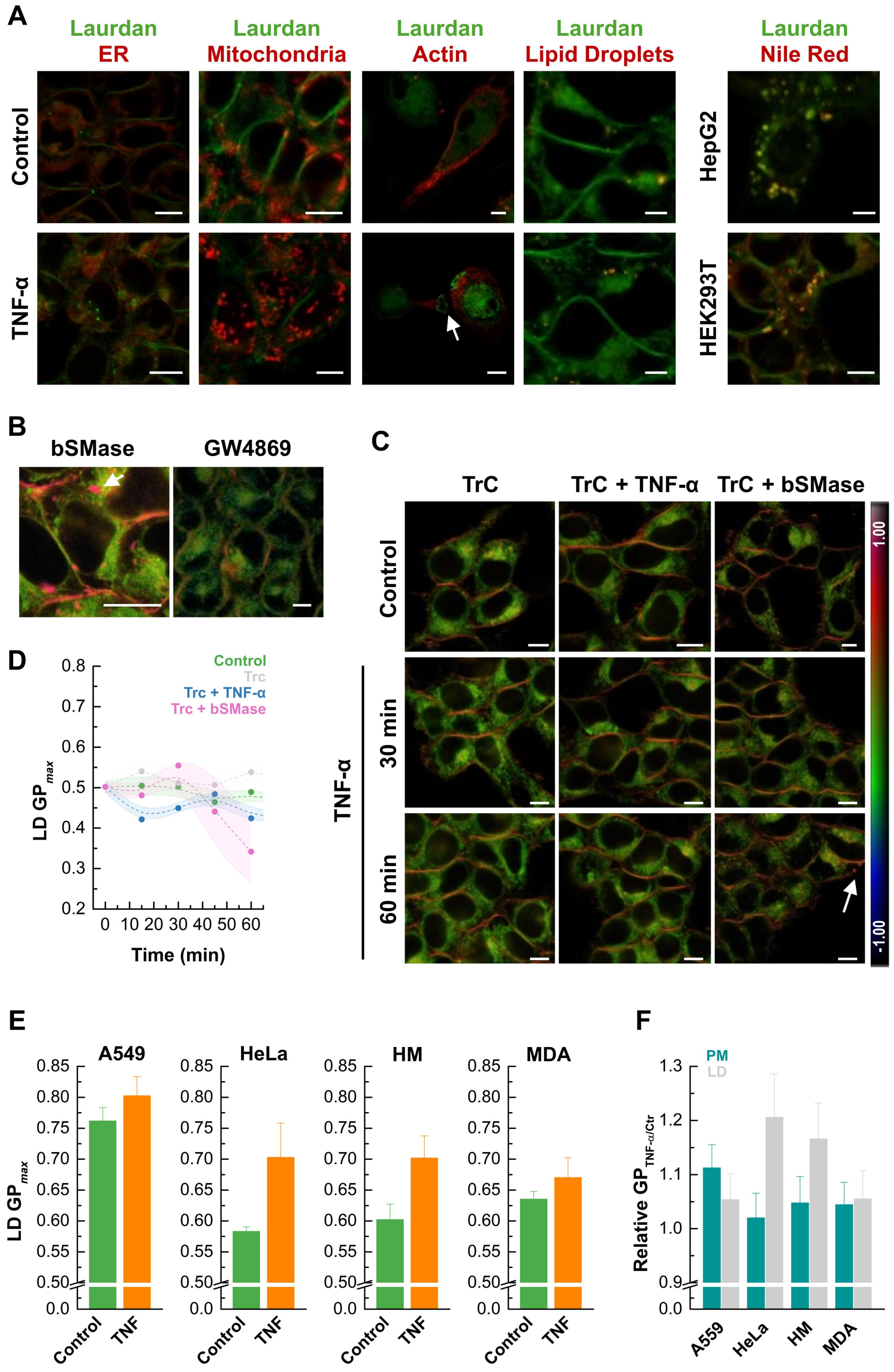


**Figure S.6. Ceramide-dependent alterations in lipid droplet properties. A)** Colocalization of Laurdan signal with organelle markers in control and TNF-α-treated cells. Cells were labeled with Laurdan and markers for the endoplasmic reticulum (ER-Tracker Red), mitochondria (MitoTracker Red FM), actin (Rhodamine-Phalloidin), and lipid droplets (LipidSpot and Nile Red). Laurdan emission from channel 1 (400–460 nm) is overlaid with organelle-specific signals. **B, C)** Representative Laurdan GP color-coded images of cells treated with **B)** bSMase or GW4869 and **C)** TrC in the absence or presence of TNF-α or bSMase. Scale bar 15 μm. **D)** Inhibition of neutral lipid synthesis by TrC reduced LDs Laurdan GPmax values. Shaded areas represent SD of at least three independent experiments. Dashed lines are included solely to guide the eye and do not represent curve fitting. **E)** Quantification of LD Laurdan GP$_{max}$ values across cell lines demonstrates ceramide-dependent modulation of LD. **F)** Relative changes in PM properties (turquoise) and LDs polarity (grey) in cells stimulated with TNF-α compared to control cells.

**Table S. 1. Kinetic parameters from one-phase exponential fits of experimental measurements in cells treated with bSMase.** Experimental data were fit to a one-phase exponential association or decay model of the form $y = A \times e^{(x/t)} + y_o$, where A is the amplitude, *t* the time constant, and $y_o$ the offset. The table presents the best-fit parameters with their associated standard deviations (± SD). The rate constant (k = 1/t) is derived from the fit. The correlation coefficient ($R^2$) indicates the goodness of fit. Comparisons include NBD and FM4-64 internalization kinetics, t-PnA anisotropy, ceramide formation, ratio of the Bodipy fluorescence intensity at 507 and 613 nm, and Pearson correlation coefficients (PCC) obtained for early and late endosomal/lysosomal compartments.

| Figure | Comparison | y0 (± SD) | A (± SD) | t (± SD) | k (1/ t) | $R^2$ |
|---|---|---|---|---|---|---|
| **S3D** | *FM4-64 Internalization vs Time* | 5.89 ± 0.82 | -4.92 ± 0.82 | -25.95 ± 8.29 | -0.04 ± 0.01 | 0.97 |
| **S3D** | *NBD Internalization vs Time* | 78.18 ± 4.30 | -55.84 ± 4.47 | -15.51 ± 3.66 | -0.06 ± 0.01 | 0.98 |
| **S3F** | *NBD Internalization vs FM4-64 Internalization* | 91.37 ± 4.74 | -93.69 ± 2.44 | -3.06 ± 0.43 | -0.33 ± 0.05 | 1.00 |
| **S3E** | *NBD Internalization vs Ceramide Formation* | 23.46 ± 3.01 | 0.002 ± 0.003 | 0.08 ± 0.02 | 11.84 ± 2.44 | 0.97 |
| **S4B** | *PCC vs NBD Internalization (EE)* | -0.59 ± 0.14 | -0.71 ± 0.14 | -24.86 ± 9.44 | -0.04 ± 0.01 | 0.96 |
| **S4B** | *PCC vs NBD Internalization (LE/Lys)* | 1.29 ± 1.36 | -1.37 ± 1.33 | -97.95 ± 134.17 | -0.01 ± 0.01 | 0.96 |
